# Early dynamics of a neutrophil interferon program drive immunotherapy resistance and limit durable anti-tumor immunity

**DOI:** 10.64898/2026.09.30.755684

**Authors:** Yang Gao, Emmanuel Anning, Linhua Wang, Yating Cheng, Ling Wu, Jun Wang, Fengshuo Liu, Zhan Xu, Xiaoxin Hao, Jun Liu, Liqun Yu, Yunfeng Ding, Weijie Zhang, David G. Edwards, Yi-Hsuan Wu, Hilda L. Chan, Dane Hoffman, Siyue Wang, Tobie D. Lee, Luis B. Dominguez, Noa Peled, Xuan Li, Xiang Chen, Charlotte H. Rivas, Na Zhao, Sebastian J. Calderon, Diego A. Pedroza, Alex J. Smith, Nan Guan, Zhandong Liu, Jeffrey M. Rosen, Xiang H.-F. Zhang

## Abstract

Despite the remarkable success of immune checkpoint blockade (ICB), durable clinical benefit remains limited to a subset of patients, highlighting the need for predictive biomarkers and new therapeutic targets. Here, we investigated the early immune dynamics associated with anti-PD1 responsiveness using longitudinal single-cell transcriptomics, spatial transcriptomics, and neutrophil-specific genetic perturbations in preclinical models of triple-negative breast cancer. We identified a conserved neutrophil interferon-stimulated gene (ISG) program that emerges before overt tumor divergence and predicts resistance to anti-PD1 therapy. Successful treatment was associated with an early reduction of IFN-responsive neutrophils in both blood and tumor tissues, whereas persistent neutrophil IFN signaling correlated with therapeutic resistance. Mechanistically, genetic disruption of either type I or type II interferon signaling in neutrophils enhanced anti-PD1 efficacy and promoted durable anti-tumor immune memory. Spatial analyses further revealed an inverse association between IFN-responsive neutrophils and tertiary lymphoid structure-enriched regions during effective therapy. Together, these findings identify IFN-responsive neutrophils as early determinants of immunotherapy outcome and establish dynamic neutrophil interferon programs as candidate blood-based biomarkers and therapeutic targets for cancer immunotherapy.

## Introduction

The advent of immune checkpoint blockade (ICB) therapies, particularly anti-PD1/PD-L1 and anti-CTLA4 antibodies, has revolutionized the treatment of various cancers by harnessing the immune system to combat malignancies.^1^ However, durable clinical benefit remains limited to a subset of patients, and both primary and acquired resistance continue to represent major clinical challenges.^2,3^ This challenge is particularly evident in triple-negative breast cancer (TNBC), a disease characterized by substantial molecular and immune heterogeneity and variable responsiveness to ICB.^4–7^ Defining the early immune events that distinguish responders from non-responders before overt tumor regression occurs is therefore essential for developing predictive biomarkers and identifying new therapeutic strategies.

The tumor microenvironment is a major determinant of ICB response.^8^ Although cytotoxic T cells, dendritic cells, and tertiary lymphoid structures (TLSs) are widely recognized as positive correlates of anti-tumor immunity,^9–11^ myeloid cells can profoundly influence whether these immune programs are effectively initiated and sustained.^12^ Neutrophils are particularly important in this context.^13–15^ Elevated peripheral neutrophil abundance, increased neutrophil-to-lymphocyte ratio, and tumor-associated neutrophil infiltration are often associated with poor outcomes and resistance to ICB.^15–17^ Consistent with these observations, our previous studies identified a neutrophil-enriched subtype of TNBC characterized by systemic neutrophilia, extensive tumor-infiltrating neutrophils, and profound resistance to checkpoint blockade.^18^ Together, these findings implicate neutrophils as important regulators of immunotherapy response and raise the possibility that specific neutrophil states may influence therapeutic outcome.

Recent single-cell studies have revealed that neutrophils comprise diverse transcriptional and functional states that can either promote or restrain anti-tumor immunity.^19–21^ HLA-DR^+^ neutrophil populations exhibit antigen-presenting properties and contribute to T cell-mediated tumor control,^19^ whereas CCL4^+^ and PD-L1^+^ neutrophils have been linked to immunosuppressive programs associated with tumor progression and therapeutic resistance.^20^ Notably, two recent studies have identified interferon-responsive neutrophil populations associated with favorable responses to immunotherapy.^22,23^ Therapy-induced IFN-responsive neutrophils, including Ly6E^hi^ or IRF1^+^ neutrophils, emerge during successful responses to ICB, support anti-tumor immune responses, and/or serve as blood-borne predictors of ICB efficacy.^22,23^ Together, these findings challenge the traditional view of tumor-associated neutrophils as predominantly immunosuppressive cells and suggest that specific neutrophil states may actively contribute to tumor rejection.

However, emerging evidence indicates that interferon signaling in neutrophils can also be immunosuppressive and promote immunotherapy resistance. A distinct IFIT1^+^ISG15^+^ neutrophil state has been associated with elevated expression of PD-L1 and features of immune suppression.^19^ Likewise, a recent study demonstrated that immunotherapy-induced IFN-gamma (IFNγ) drives neutrophils toward an aged, PD-L1-expressing, immunosuppressive phenotype that limits treatment efficacy.^24^ In that setting, neutrophil-specific disruption of IFNγ receptor signaling prevented the development of suppressive neutrophils and enhanced anti-tumor responses.^24^ Together, these findings suggest that the functional consequences of interferon signaling in neutrophils may be highly context dependent. Whether distinct interferon-responsive neutrophil states promote or restrain anti-tumor immunity, and how these divergent outcomes are determined during immunotherapy, remain unresolved questions.

Here, we investigated the early immune dynamics associated with anti-PD1 responsiveness in preclinical TNBC models displaying heterogeneous therapeutic outcomes. By integrating longitudinal single-cell transcriptomic profiling, spatial transcriptomics, and neutrophil-specific genetic perturbations, we identified a neutrophil-restricted interferon-stimulated gene program that predicts and promotes resistance to checkpoint blockade. Our findings identify IFN-responsive neutrophils as early determinants of anti-PD1 efficacy and establish dynamic neutrophil interferon programs as candidate biomarkers and therapeutic targets for cancer immunotherapy.

## Results

### Early phase decrease in blood neutrophil interferon responses predicts anti-PD1 therapy efficacy in preclinical mouse models of breast cancer

To investigate the potential determinants of immunotherapy efficacy, we examined a few syngeneic murine TNBC models that were previously characterized by the tumor immune microenvironment enriched with macrophages but low in neutrophils.^18^ We focused on these models because of their intragroup heterogeneous responses to the anti-PD1 treatment, i.e., different biological replicates among the same treatment group can exhibit either complete regression or almost unaffected growth (**Figure 1A – 1C**). Interestingly, the scarcity of neutrophils seems to be a pre-requisite of the observed intragroup dichotomy because none of the biological replicates responded to anti-PD1 in several neutrophil-enriched tumor models (**Figure S1A – S1G**). The dichotomous response in the neutrophil-poor models is remarkable because it occurred despite the common cancer model, the same treatment regimen, and the genetically identical hosts. To understand this better, we designed experiments to identify molecular and cellular events that may distinguish responders from non-responders. We reasoned that these events can only be delineated before evident tumor regression, as the latter will be confounded by dramatic cell death and inflammatory responses as consequences but not causes of successful anti-tumor immunity.

**Figure 1.**
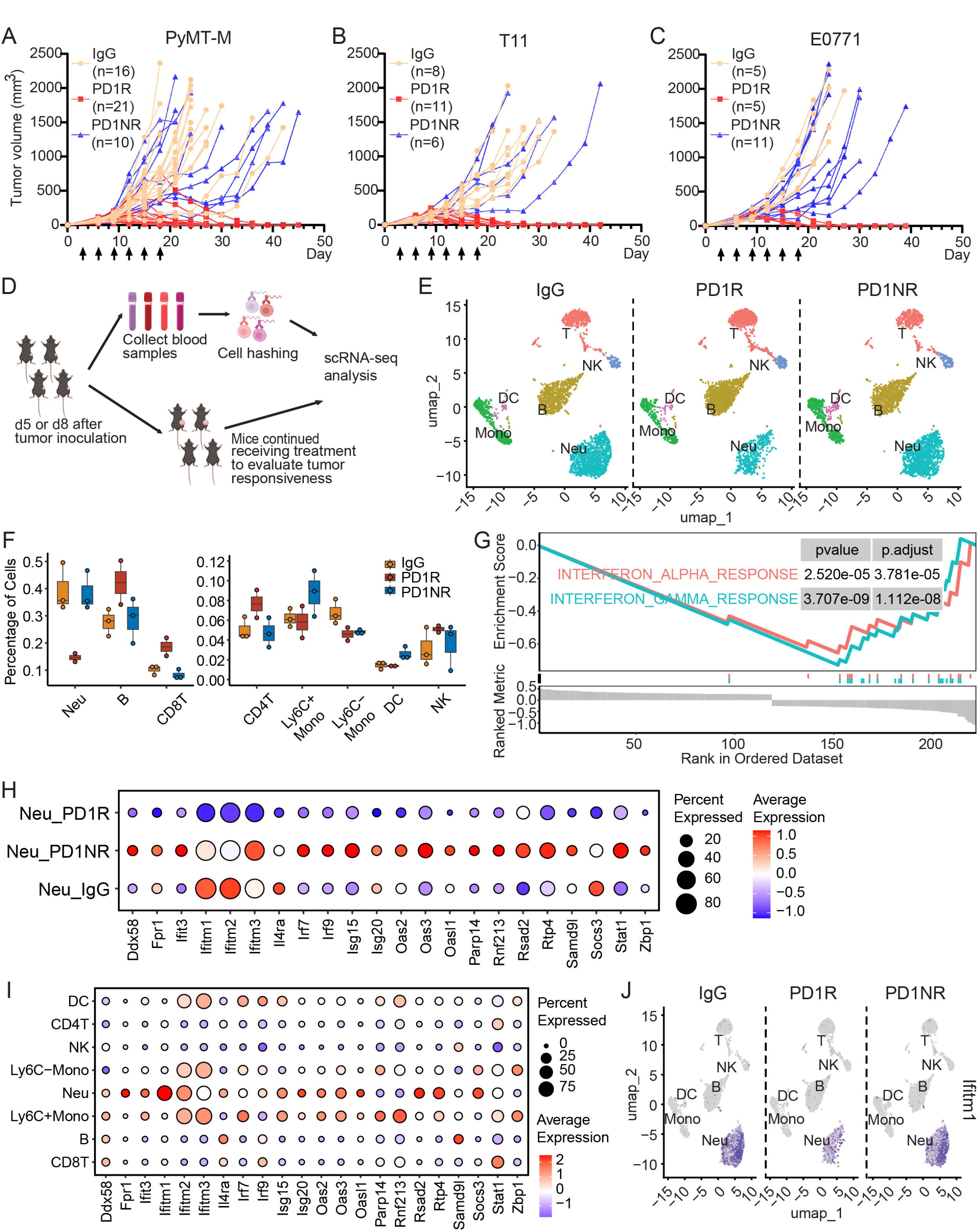
Early neutrophil interferon responses distinguish responders from non-responders before tumor regression. **(A–C)** Tumor growth curves demonstrating heterogeneous responses to anti-PD1 therapy in three syngeneic TNBC models. Individual mice were classified as responders (PD1R) or non-responders (PD1NR) based on final therapeutic outcome. Arrows indicate anti-PD1 antibody administration. **(D)** Experimental workflow for longitudinal immune profiling. Peripheral blood was collected on day 5 or day 8 following tumor inoculation, prior to overt tumor regression, for cell hashing-based single-cell RNA sequencing (scRNA-seq). Mice subsequently continued anti-PD1 treatment to determine therapeutic responsiveness. **(E)** UMAP visualization of peripheral blood immune cells from IgG-treated controls, anti-PD1 responders (PD1R), and anti-PD1 non-responders (PD1NR), identifying major immune cell populations. **(F)** Quantification of immune cell populations in peripheral blood. Responders exhibited increased B cells and T cells together with decreased neutrophils compared with non-responders. **(G)** Gene set enrichment analysis (GSEA) of neutrophils comparing responders and non-responders revealed significant negative enrichment of interferon-α and interferon-γ response pathways in responders. Normalized enrichment scores (NES), *P* values, and adjusted *P* values are shown. **(H)** Dot plot showing expression of 22 interferon-stimulated genes (ISGs) in neutrophils from IgG-treated controls, responders, and non-responders. **(I)** Dot plot illustrating expression of the identified ISGs across peripheral blood immune cell populations. Ifitm1 exhibited the greatest neutrophil specificity among the identified ISGs. **(J)** Feature plots showing Ifitm1 expression across peripheral blood immune cell populations from IgG-treated controls, responders, and non-responders, demonstrating predominant expression in neutrophils. N = 3 for IgG-treated controls, n = 2 for PD1R, and n = 3 for PD1NR.

We first set out to monitor immune alterations that may precede tumor regression and be reflected in the peripheral blood.^25^ To this end, we collected blood samples at early time points after treatment, when tumor sizes were comparable between responders and non-responders. The immune cells from these samples were then subjected to cell hashing and single-cell RNA sequencing (scRNA-seq). Meanwhile, the mice continued to receive treatment and were inspected to evaluate tumor responsiveness (**Figure 1D**). A comparison of PyMT-M blood samples collected on day 5 (i.e., five days post tumor inoculation and one dose of anti-PD1 antibody treatment) did not reveal significant differences in immune cell populations between responders and non-responders (**Figure S1H and S1I**). However, by day 8 (i.e., eight days post tumor inoculation and two doses of anti-PD1 antibody treatment), an increase in B cells and CD8+ T cells, along with a decrease in neutrophils, was observed (**Figure 1E and 1F**). These findings align with previous studies demonstrating that an elevated neutrophil-to-lymphocyte ratio is associated with poorer outcomes following ICB therapy across all major cancer types, including breast cancer.^16,17^

To identify molecular pathways associated with anti-PD1 therapy response, we performed gene set enrichment analysis (GSEA) comparing the single-cell transcriptomic profiles of responders and non-responders in PyMT-M Day 8 samples. Notably, no significantly enriched hallmark pathways were identified in B cells, T cells, or monocytes. In contrast, neutrophils displayed significant negative enrichment of both interferon-alpha (IFN-α) and IFN-γ response pathways in responders (**Figure 1G**). Among the 22 significantly altered IFN-stimulated genes (ISGs) on Day 8 (**Figure 1H**), 16 were also downregulated in responders compared to non-responders on Day 5, although only *Ifitm1* and *Ifitm3* reached statistically significance at this earlier time point (**Figure S1J**). These findings identify a conserved neutrophil ISG program that emerges prior to overt tumor regression and is inversely associated with anti-PD1 efficacy, suggesting that early neutrophil interferon responses may influence subsequent therapeutic outcome.

To identify neutrophil-specific ISGs as potential biomarkers, we evaluated the expression of these 22 ISGs across various immune cell types in the blood. Among these, *Ifitm1* emerged as the most neutrophil-specific ISG, followed by *Fpr1*, *Rsad2*, *Oasl1*, *Isg20*, and *Socs3* to a lesser extent, consistently across PyMT-M-day-8 and PyMT-M-day-5 blood samples (**Figure 1I, 1J, and S1K**). These findings suggest that neutrophil-specific ISGs, particularly *Ifitm1*, may serve as promising biomarkers for predicting responsiveness to anti-PD1 therapy.

### Early-Phase tumor samples mirror immune cell dynamics observed in blood

Next, we aimed to investigate the immune cell dynamics within tumor tissues during anti-PD1 treatment. Again, we used an approach that allowed us to capture molecular and cellular changes preceding regression.^26,27^ Specifically, we adopted a bilateral tumor inoculation model, which demonstrates a systemic and symmetrical response to both anti-IgG and anti-PD1 therapy (**Figure 2A**). We collected tumor samples from the left flank at day 8 for 10X Visium spatial transcriptomic analysis. Meanwhile, treatment was continued on the contralateral tumors to assess therapy responsiveness over time (**Figure 2B**). This approach enabled us to link early immune responses in the tumor microenvironment to subsequent treatment outcomes, providing insights into the mechanisms driving differential responses to anti-PD1 therapy. Spatial transcriptomic deconvolution using Robust Cell Type Decomposition (RCTD) identified consistent immune cell dynamics between tumor and blood samples, including an increase in B cells and T cells and a decrease in neutrophils within tumor tissues of responders (**Figure 2C and 2D**). These findings were corroborated by an independent deconvolution method, cell2location, which revealed similar patterns of immune cell infiltration and abundance (**Figure S2A and S2B**). Comparisons of Gene Ontology (GO) Terms between responder tumor samples and those from non-responders or controls highlighted significant enrichment in pathways associated with B cell and T cell activation, proliferation, and differentiation in responders (**Figure 2E – 2G**).

**Figure 2.**
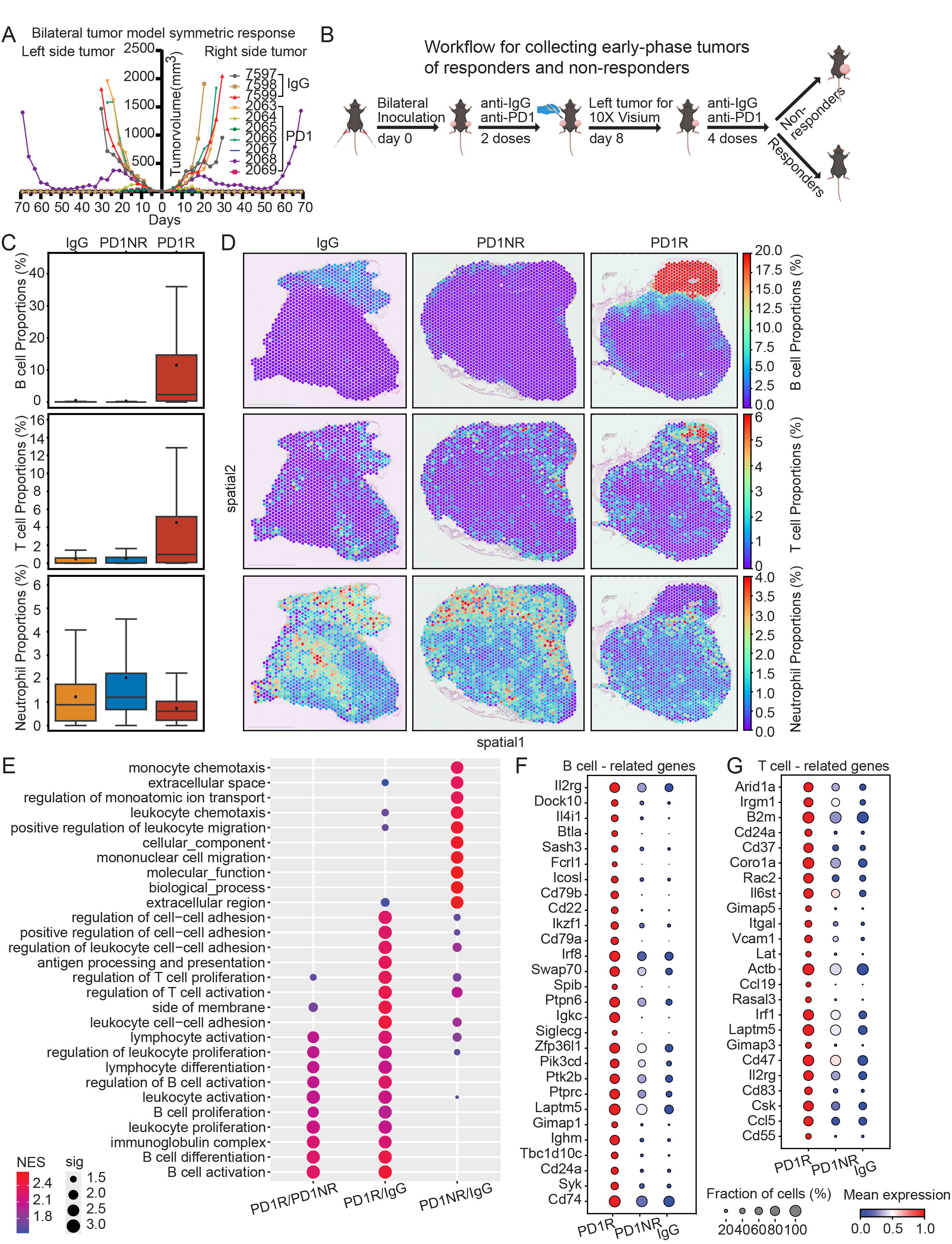
Early tumor immune remodeling mirrors peripheral blood immune dynamics during anti-PD1 therapy. **(A)** Bilateral tumor model demonstrating symmetric responses to anti-PD1 therapy. Mice were inoculated with tumors on both side of the 4^th^ mammary fat pads, and tumor growth on the left and right sides showed highly concordant responses to anti-PD1 treatment. **(B)** Experimental workflow for early-phase spatial transcriptomic analysis. Mice bearing bilateral tumors received anti-IgG or anti-PD1 antibodies. One tumor was harvested on day 8 for 10X Visium spatial transcriptomic analysis before overt tumor regression, while treatment continued on the contralateral tumor to classify mice as responders (PD1R) or non-responders (PD1NR). **(C)** Cell-type deconvolution of Visium spatial transcriptomic data showing increased proportions of B cells and T cells, together with reduced neutrophils, in responder tumors compared with non-responder and IgG-treated tumors. **(D)** Representative spatial distribution maps of B cells, T cells, and neutrophils generated by RCTD deconvolution across IgG-treated, PD1NR, and PD1R tumors. **(E)** Gene Ontology (GO) enrichment analysis of differentially expressed genes in responder tumors. Compared with non-responders or IgG-treated controls, responder tumors were enriched for biological processes associated with B-cell activation, lymphocyte activation and differentiation, antigen processing and presentation, leukocyte proliferation, and cell-cell adhesion. **(F)** Dot plot showing expression of representative B cell-associated genes in spatial transcriptomic data across IgG-treated, PD1NR, and PD1R tumors. **(G)** Dot plot showing expression of representative T cell-associated genes in spatial transcriptomic data across IgG-treated, PD1NR, and PD1R tumors. N = 2 per group for IgG-treated control, PD1R, and PD1NR tumors.

### Interferon-responsive neutrophils drive tumors resistant to anti-PD1 therapy

Further analysis of pairwise cell types using Spearman’s correlation revealed a strong spatial co-localization among B cells, T cells, and dendritic cells, suggesting the presence of TLSs in responder tumors (**Figure 3A**).^28,29^ This finding was corroborated by immunofluorescence staining, which confirmed the presence of organized immune cell aggregates characteristic of TLSs (**Figure 3B**). While previous preclinical and clinical studies have largely focused on lymphocyte-associated biomarkers of immunotherapy response,^28,30–34^ our prior work demonstrated that neutrophils can act as potent mediators of resistance in neutrophil-enriched tumors.^18^ In the present study, we focused on tumors initially characterized by low neutrophil abundance. Remarkably, the small population of residual neutrophils became further excluded from lymphocyte-rich regions during successful anti-PD1 therapy (**Figure 3A**), suggesting that spatial segregation of neutrophils from TLS-associated immune niches may represent an early hallmark of therapeutic responsiveness.

**Figure 3.**
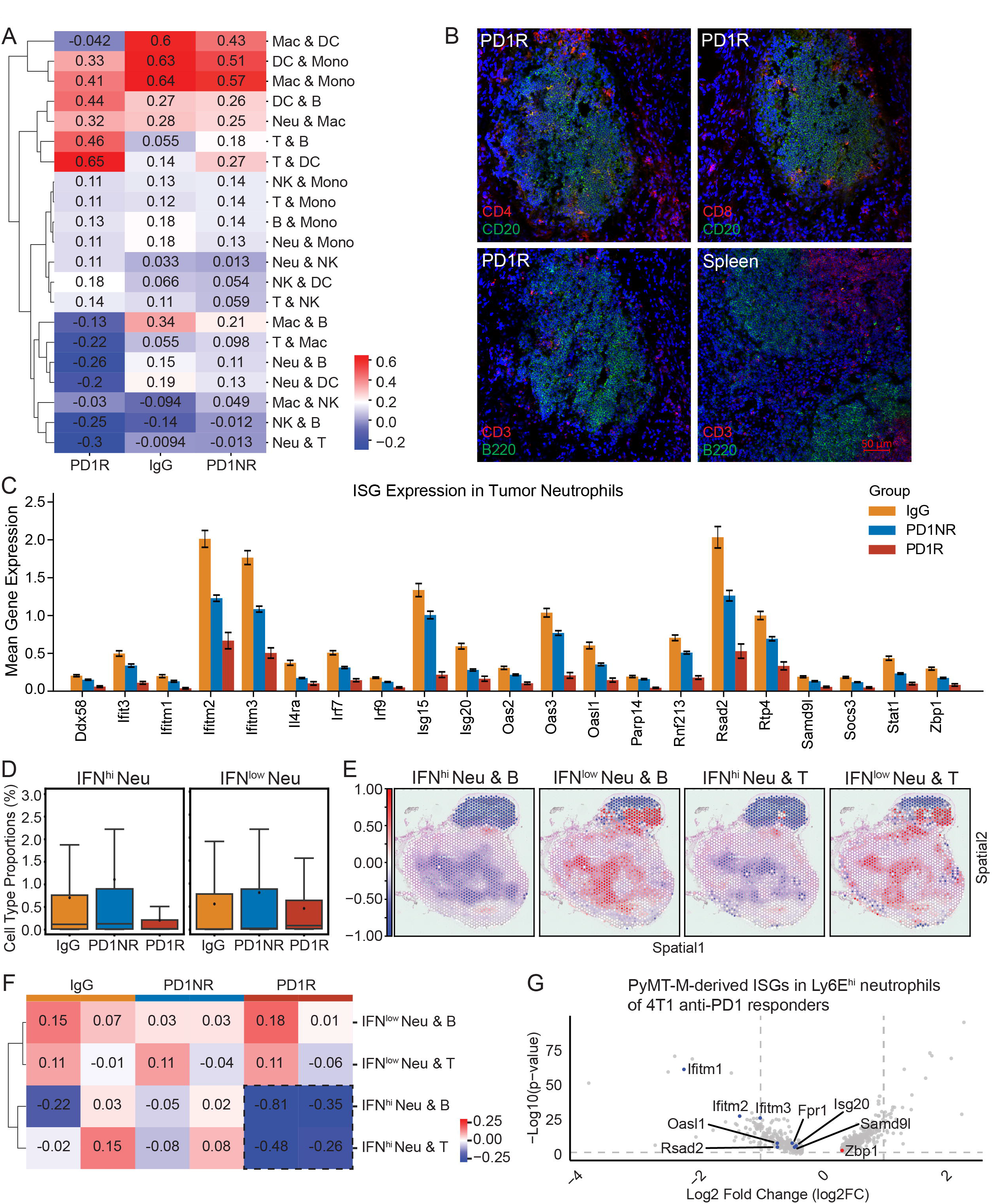
IFN-responsive neutrophils are spatially excluded from TLS-associated immune niches during successful anti-PD1 therapy. **(A)** Spatial correlation analysis of immune cell populations in IgG-treated controls, anti-PD1 non-responders (PD1NR), and anti-PD1 responders (PD1R). Spearman correlation coefficients were calculated from cell-type proportions inferred by spatial transcriptomic deconvolution. Responding tumors exhibited increased spatial association among B cells, T cells, and dendritic cells, consistent with the formation of tertiary lymphoid structure (TLS)-associated immune niches, while neutrophils displayed reduced association with lymphocyte populations. **(B)** Representative immunofluorescence images showing TLS-like immune aggregates in responder tumors. Tumor sections were stained for CD4, CD8, CD3, CD20, and B220. Mouse spleen served as a positive control. Scale bar, 50 μm. **(C)** Mean expression of the neutrophil interferon-stimulated genes (ISGs) identified in peripheral blood (Figure 1) in tumor-associated neutrophils from IgG-treated controls, PD1NR, and PD1R tumors. Expression of multiple ISGs was significantly reduced in responder tumors. **(D)** Quantification of IFN^hi^ and IFN^low^ neutrophil populations identified according to neutrophil ISG module scores using RCTD. Responding tumors contained significantly fewer IFN^hi^ neutrophils. **(E)** Representative spatial correlation maps showing the spatial relationships between IFN^hi^ or IFN^low^ neutrophils and B cells or T cells in responder tumors. IFN^hi^ neutrophils exhibited markedly reduced spatial association with lymphocyte-rich regions compared with IFN^low^ neutrophils. **(F)** Heatmap summarizing spatial correlation coefficients between IFN^hi^ or IFN^low^ neutrophils and B cells or T cells in IgG-treated controls, PD1NR, and PD1R tumors. Successful anti-PD1 therapy was associated with progressive spatial exclusion of IFN^hi^ neutrophils from TLS-associated immune niches. **(G)** Comparison of the PyMT-M IFN-responsive neutrophil signature with Ly6E^hi^ neutrophils from anti-PD1-responsive 4T1 tumors reported by Benguigui *et al.* Volcano plot highlighting the expression of PyMT-M-derived ISGs in Ly6E^hi^ neutrophils. Several ISGs associated with resistant neutrophils in the present study, including members of the *Ifitm* family, *Oasl1*, *Rsad2*, and *Samd9l*, were downregulated in Ly6E^hi^ neutrophils, suggesting that distinct interferon-associated neutrophil programs underlie divergent immunotherapy outcomes. N = 2 per group for IgG-treated control, PD1R, and PD1NR tumors.

Next, we analyzed the expression of ISGs in tumor-infiltrating neutrophils across different treatment groups. Consistent with our observations from circulating neutrophils in peripheral blood, all 22 ISGs exhibited significant downregulation in tumor-infiltrating neutrophils from responders (**Figure 3C**). Consistent with our observations in circulating peripheral blood neutrophils, all 22 ISGs exhibited significant downregulation in tumor-infiltrating neutrophils from responders. Notably, Ifitm1 again emerged as a neutrophil-specific ISG within the tumor microenvironment, further highlighting its potential utility as a predictive biomarker for anti-PD1 therapy response (**Figure S3A**). These results reinforce the critical role of neutrophil-specific IFN signaling pathways in mediating resistance to immunotherapy.

We also stratified neutrophils into IFN^hi^ and IFN^low^ populations based on the expression of the identified ISG signature. Notably, IFN^hi^ neutrophils, but not IFN^low^ neutrophils, were dramatically reduced in responder tumor samples during the early phase of anti-PD1 therapy (**Figure 3D and S3B**). Furthermore, IFN^hi^ neutrophils exhibited spatial exclusion from regions enriched with B and T cells, particularly within TLSs, in responder tumors (**Figure 3E**), and this exclusion was quantitatively assessed by our in-house-developed algorithm, Spatial Enrichment Analysis of Gene Associations using *L*-index (SEAGAL)^35^ (**Figure 3F**), and independently validated using commonly used correlation-based approaches, including Pearson’s correlation (**Figure S3C**), and Spearman’s correlation (**Figure S3D**). Taken together, these findings suggest that neutrophils, especially the IFN^hi^ subset, are excluded from regions enriched with lymphocytes or TLSs.

We then sought to validate our findings using additional models to further investigate the role of IFN-responsive neutrophils in anti-PD1 therapy. Since we observed the strong *de novo* resistance of models with high baseline level of neutrophils, we compared neutrophils from these models to those derived from models with initially low neutrophil frequency. We paired the models based on their genetic background and the genetic drivers (**Figure S1C – S1G**).

Specifically, PyMT-N is an isogenic sister cell line of PyMT-M, whereas 2208L and T11 are both derived P53KO Balb/c mice^18^. PyMT-N and 2208L models both enrich neutrophils locally and systemically compared to PyMT-M and T11 according to our previous studies.^18^ Consistent with our observations, 2208L neutrophils showed enrichment in the IFN response pathways (**Figure S3E**). At first glance, these findings appear inconsistent with a recent study reporting that IFN-stimulated Ly6E^hi^ neutrophils serve as favorable prognostic biomarkers for anti-PD1 therapy response in 4T1 models.^23^ However, a closer examination revealed that Ly6E^hi^ neutrophils also exhibit downregulation of several ISGs — including *Ifitm1*, *Ifitm2*, and *Ifitm3* — which were similarly downregulated in responders in our PyMT-M model (**Figure 3G**). Collectively, these observations suggest that distinct neutrophil ISGs may exert divergent effects on anti-tumor immunity.

Interestingly, cancer cells from models with high neutrophil abundance also exhibited a marked upregulation of both IFN-α and IFN-γ pathways compared to their paired counterparts, suggesting that tumor-intrinsic IFN signaling may be linked to the increased expression of ISGs in the neutrophils (**Figure S3F**). To validate this observation, we performed *in vitro* coculture experiments using tumor cells and naïve bone marrow-derived neutrophils. Indeed, PyMT-N tumor cells elicited a stronger upregulation of ISGs in the cocultured neutrophils (**Figure S3G**), indicating that direct contact with tumor cells exhibiting increased IFN pathways amplifies the neutrophils’ IFN responses.

### Conditional knockout of interferon receptors in neutrophils potentiates anti-PD1 therapy response

To study the causal role of IFN-responsive neutrophils in anti-PD1 immunotherapy, we employed conditional knockout mouse models to selectively delete IFN signaling pathways in neutrophils. Notably, many ISGs can be activated by both type I and type II IFN signaling pathways, raising the possibility of compensatory effects when only one pathway is disrupted.^36–40^ Consistent with this notion, ablation of either IFNAR1 (type I IFN receptor) or IFNGR1 (type II IFN receptor) in neutrophils using the S100A8-Cre system did not significantly improve the response rate or survival of PyMT-M tumor-bearing mice (**Figure 4A–4D**).

**Figure 4.**
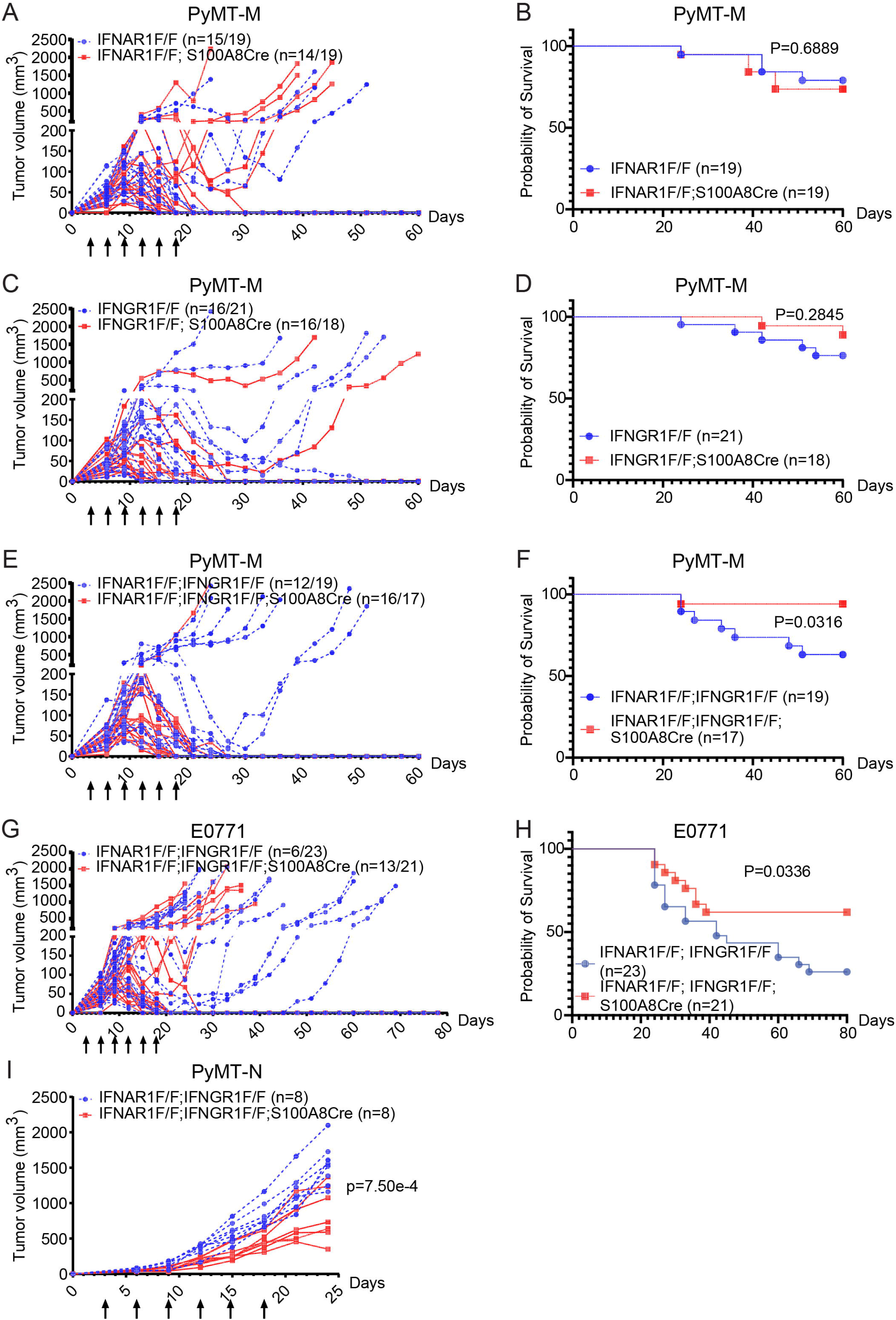
Dual inhibition of type I and type II interferon signaling in neutrophils overcomes resistance to anti-PD1 therapy. **(A, B)** Tumor growth curves (A) and Kaplan–Meier survival analysis (B) of anti-PD1-treated IFNAR1F/F and IFNAR1F/F; S100A8Cre mice bearing PyMT-M tumors. Neutrophil-specific deletion of IFNAR1 alone did not significantly improve therapeutic responses. **(C, D)** Tumor growth curves (C) and Kaplan–Meier survival analysis (D) of anti-PD1-treated IFNGR1F/F and IFNGR1F/F; S100A8Cre mice bearing PyMT-M tumors. Neutrophil-specific deletion of IFNGR1 alone similarly failed to significantly enhance anti-PD1 efficacy. **(E, F)** Tumor growth curves (E) and Kaplan–Meier survival analysis (F) of anti-PD1-treated IFNAR1F/F; IFNGR1F/F and IFNAR1F/F; IFNGR1F/F; S100A8Cre mice bearing PyMT-M tumors. Simultaneous disruption of both type I and type II interferon signaling in neutrophils markedly enhanced anti-PD1 therapeutic efficacy and significantly prolonged survival. **(G, H)** Validation of dual neutrophil-specific IFNAR1/IFNGR1 deletion in the E0771 TNBC model. Tumor growth curves (G) and Kaplan–Meier survival analysis (H) demonstrate enhanced anti-PD1 responses following simultaneous disruption of both interferon signaling pathways in neutrophils. **(I)** Therapeutic responses of neutrophil-enriched, anti-PD1-resistant PyMT-N tumors following dual neutrophil-specific deletion of IFNAR1 and IFNGR1. Combined disruption of type I and type II interferon signaling significantly sensitized this intrinsically resistant model to anti-PD1 therapy. Tumor-bearing mice received anti-PD1 antibody as indicated by arrows. Individual tumor growth curves are shown. For responder frequencies, n = x/y indicates the number of PD1R mice (x) among the total number of anti-PD-1-treated mice (y). Kaplan–Meier survival curves represent the percentage of tumor-free mice over time. Statistical analyses were performed using Log-rank (Mantel-Cox) test for survival curves comparison and two-way ANOVA for tumor growth comparison. *P* values are indicated in the figure.

In contrast, simultaneous deletion of both IFNAR1 and IFNGR1 in neutrophils significantly enhanced the response rate of PyMT-M tumors to anti-PD1 therapy and effectively mitigated resistance (**Figure 4E and 4F**). Additionally, dual deletion of IFN signaling in neutrophils improved anti-PD1 therapy efficacy in E0771, another model that is initially neutrophil-poor (**Figure 4G and 4H**). Importantly, in the PyMT-N model, which is enriched with neutrophils and *de novo* resistant to anti-PD1 therapy, deletion of both IFN signaling pathways markedly slowed tumor growth (**Figure 4I**). These findings highlight the critical role of IFN signaling in neutrophil-mediated resistance to anti-PD1 therapy and suggest that targeting both type I and type II IFN pathways in neutrophils may be necessary to overcome resistance.

### IFN-responsive neutrophils impair immunological memory induced by anti-PD1 therapy

Because durable responses to checkpoint blockade are thought to depend on the generation of long-lived anti-tumor immune memory,^1,41,42^ we next investigated whether IFN-responsive neutrophils influence memory formation following successful therapy. In our models, anti-PD1 treatment clearly generated durable responses in animals without the rise of IFN-responsive neutrophils. Furthermore, E0771 tumor-bearing mice cured by ICB treatment rejected intracardiac rechallenge with E0771 cells, effectively preventing experimental metastasis, while control mice developed metastases across multiple organs (**Figure S5A and S5B**). Rechallenge with luciferase-labeled E0771 cells further confirmed that cured mice eliminated tumor cells within the first three days post-rechallenge (**Figure S5C**). Similar results were observed in PyMT-M models, where anti-PD1-cured mice rejected PyMT-M tumor cells transplanted to the contralateral side of the mammary fat pad, preventing tumor growth (**Figure 5A – 5D**). This robust immunological memory was primarily maintained by T cells, as antibody depleting T cells in cured mice restored PyMT-M tumor growth upon rechallenge (**Figure S5D**). Consistently, PyMT-M tumors showed higher levels of memory T cells infiltration compared to their isogeneic neutrophil-enriched model – PyMT-N (**Figure S5E**).

**Figure 5.**
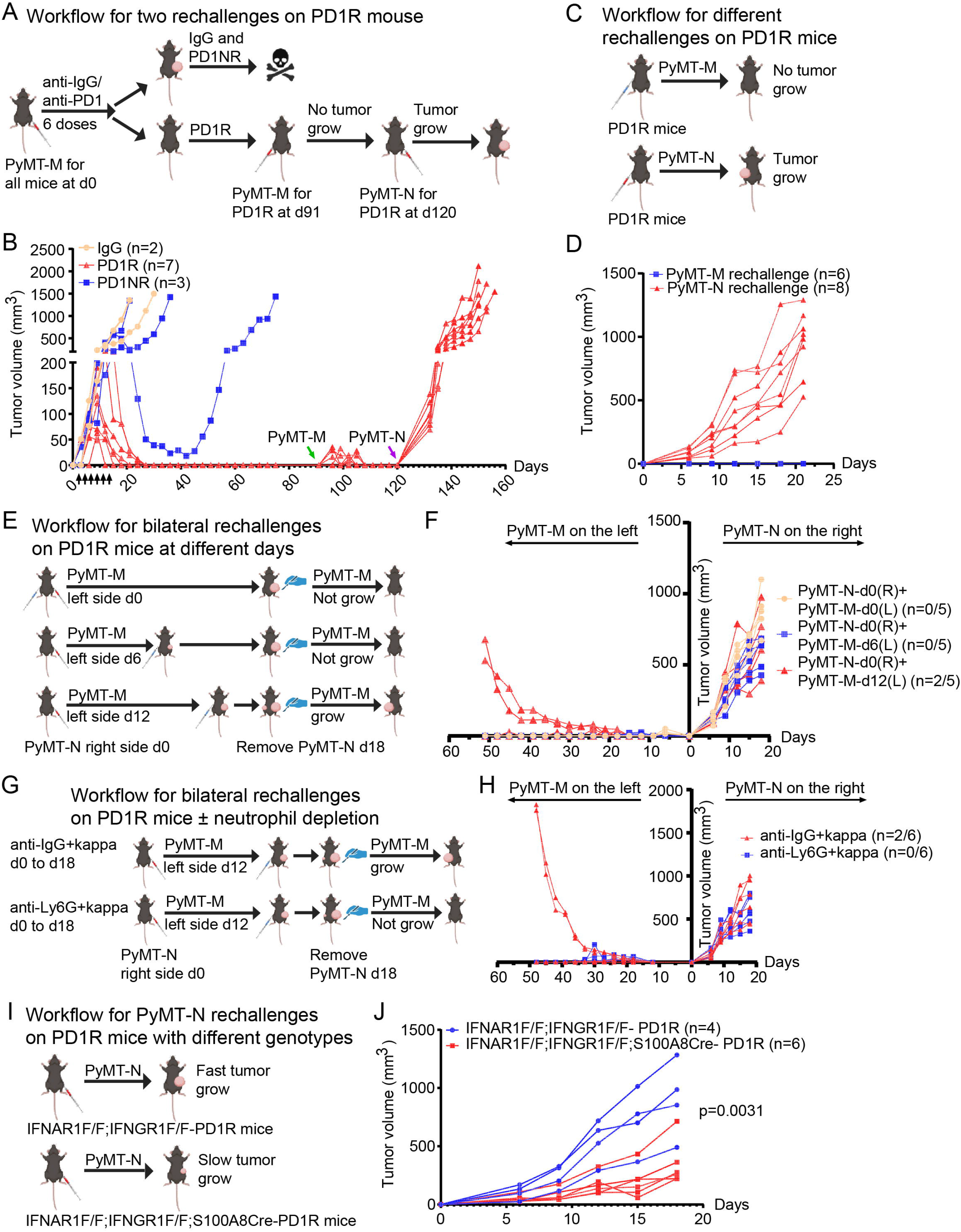
Neutrophil interferon signaling limits the establishment and maintenance of durable anti-tumor immune memory following anti-PD1 therapy. **(A)** Experimental design for sequential tumor rechallenge. Mice bearing PyMT-M tumors were treated with anti-PD1 or control IgG. Mice that completely rejected primary tumors following anti-PD1 treatment (PD1R) were rechallenged with PyMT-M tumors on day 91 and subsequently with PyMT-N tumors on day 120 to evaluate durable anti-tumor immune memory and tumor subtype specificity. **(B)** Tumor growth following sequential rechallenge. PD1R mice completely rejected PyMT-M rechallenge but failed to reject subsequent PyMT-N rechallenge. **(C)** Experimental design comparing rechallenge with either PyMT-M or PyMT-N tumors in independent cohorts of PD1R mice. **(D)** Growth kinetics of rechallenged tumors demonstrating complete protection against PyMT-M but not PyMT-N tumors, indicating tumor-specific immune memory following successful anti-PD1 therapy. **(E)** Experimental design investigating whether established PyMT-N tumors suppress anti-tumor immune memory. PD1R mice were inoculated with PyMT-N tumors on the right side, followed by delayed PyMT-M rechallenge on the contralateral side at the indicated time points. PyMT-N tumors were surgically removed on day 18. **(F)** Tumor growth following bilateral rechallenge. Early establishment of PyMT-N tumors progressively impaired rejection of contralateral PyMT-M rechallenge, indicating that neutrophil-enriched tumors suppress previously established anti-tumor immune memory. **(G)** Experimental design evaluating the contribution of neutrophils to memory suppression. PD1R mice bearing PyMT-N tumors received either control antibodies or anti-Ly6G antibody during bilateral rechallenge. **(H)** Neutrophil depletion restored rejection of contralateral PyMT-M tumors despite the presence of PyMT-N tumors, demonstrating that neutrophils are required for suppression of established anti-tumor immune memory. **(I)** Experimental design evaluating the role of neutrophil interferon signaling in suppressing immune memory. PD1R mice with or without neutrophil-specific deletion of IFNAR1/IFNGR1 were rechallenged with PyMT-N tumors. **(J)** PyMT-N tumor growth following rechallenge in control and neutrophil-specific IFNAR/IFNGR1 double-knockout PD1R mice. Dual deletion of IFNAR1 and IFNGR1 in neutrophils significantly reduced PyMT-N tumor outgrowth following rechallenge, indicating that neutrophil interferon signaling contributes to suppression of established anti-tumor immune memory. Data are presented as individual biological replicates. For tumor rechallenge experiments, n = x/y indicates the number of mice with PyMT-M tumor growth (x) among the total number of rechallenged mice (y). Statistical analyses were performed using two-way ANOVA. *P* values are indicated in the figure.

The immunological memory in cured mice failed to prevent tumor growth when rechallenged with PyMT-N (**Figure 5A – 5D**). This may not be surprising given that PyMT-M and PyMT-N, although derived from the same tumor, may express divergent antigens. However, when we implanted PyMT-N tumor cells to PyMT-M-cured mice, the immune memory against PyMT-M cells appeared to be impaired because 2 of 5 mice now developed tumors on the contralateral side when they were rechallenged with the PyMT-M cells (**Figure 5E and 5F**).

To determine whether neutrophils mediated this suppression, we performed neutrophil depletion using anti-Ly6G and anti-rat kappa light chain antibodies. Indeed, this depletion restored the ability of hosts to reject PyMT-M rechallenge despite the presence of PyMT-N tumors (**Figure 5G and 5H**). These findings suggest that neutrophils mediate the inhibition of immunological memory in the cured mice by PyMT-N tumors.

Furthermore, we examined whether neutrophil IFN signaling played a role in suppressing immunological memory by using cured mice with dual knockout of IFNAR1 and IFNGR1. Interestingly, loss of IFN signaling suppressed PyMT-N tumor growth in the cured mice, indicating that IFN-responsive neutrophils partially mediate the inhibition of immunological memory induced by PyMT-N (**Figure 5I and 5J**). Together, these findings demonstrate that neutrophil-intrinsic interferon signaling limits the establishment and maintenance of durable anti-tumor immune memory.

## Discussion

Immune checkpoint blockade can induce durable tumor regression, yet the early immune events that determine whether therapy succeeds or fails remain incompletely understood.^43,44^ Here, we identify an early neutrophil interferon program that predicts and functionally contributes to resistance to anti-PD1 therapy in preclinical models of triple-negative breast cancer. By integrating longitudinal single-cell transcriptomics of peripheral blood, spatial transcriptomics of tumor tissues, and neutrophil-specific genetic perturbations, we demonstrate that successful immunotherapy is accompanied by rapid attenuation of a neutrophil-restricted ISG program before overt tumor regression. Conversely, persistence of this program is associated with therapeutic resistance. Furthermore, combined disruption of type I and type II interferon signaling in neutrophils enhances anti-PD1 efficacy and promotes durable anti-tumor immune memory. Together, these findings establish dynamic neutrophil interferon programs as early determinants of immunotherapy outcome and identify them as candidate biomarkers and therapeutic targets.

A major strength of this study is the emphasis on immune dynamics before measurable tumor regression. Most previous investigations have compared tumors after responders and non-responders have already diverged, making it difficult to distinguish causal immune programs from secondary consequences of tumor destruction and tissue remodeling. By interrogating both blood and tumor tissues while tumor burdens remained comparable, we identified a neutrophil-restricted interferon program that consistently preceded therapeutic divergence. Importantly, this program was readily detectable in peripheral blood, suggesting that longitudinal monitoring of circulating neutrophil states may provide predictive information substantially earlier than conventional radiographic assessment or endpoint tissue analyses. These findings support the concept that dynamic systemic immune profiling, rather than static endpoint measurements, may offer greater insight into the mechanisms governing immunotherapy response.

The prominence of interferon signaling in our study was somewhat unexpected given the well-established anti-tumor functions of both type I and type II interferons.^36^ These cytokines enhance antigen presentation, dendritic cell activation, T-cell priming, and cytotoxic effector function, making them central regulators of anti-tumor immunity.^40^ At the same time, persistent or dysregulated interferon signaling can induce immune-evasion programs in tumor cells and immunosuppressive phenotypes in myeloid populations,^45,46^ illustrating the pleiotropic nature of interferon biology during cancer therapy. Consequently, the functional significance of interferon signaling in neutrophils has remained incompletely understood.

Recent single-cell studies further highlight this complexity. Benguigui and colleagues identified Ly6E^hi^ interferon-responsive neutrophils that accumulate during successful anti-PD1 therapy and promote anti-tumor immunity,^23^ whereas Hirschhorn et al. demonstrated that IRF1^+^ neutrophils contribute to T cell-mediated tumor elimination.^21^ In contrast, Wu et al. identified an IFIT1^+^ISG15^+^ interferon-stimulated neutrophil population characterized by elevated PD-L1 expression and immunosuppressive features,^19^ while Pei et al. showed that immunotherapy-induced IFNγ signaling drives neutrophils toward an aged, PD-L1-expressing phenotype that limits checkpoint blockade efficacy.^24^ These observations appear contradictory and raise the fundamental question of whether interferon-responsive neutrophils are beneficial or detrimental during cancer immunotherapy, and a framework for reconciling these seemingly divergent observations is heavily needed.

On top of these previous studies, our data indicate that the temporal dynamics of these interferon programs are equally informative. By interrogating immune responses before measurable tumor regression, we found that persistence of a neutrophil-restricted ISG program early during therapy consistently marked subsequent resistance, whereas successful anti-PD1 treatment was accompanied by rapid attenuation of this program. Moreover, comparison with previously reported Ly6E^hi^ neutrophils revealed that several ISGs defining our resistant neutrophil program, including members of the Ifitm family, were downregulated in the Ly6E^hi^ population, suggesting that distinct interferon-associated neutrophil states differ not only in abundance but also in the composition of their downstream transcriptional programs. Collectively, these observations support a model in which the biological function of neutrophil interferon signaling is determined not simply by pathway activation, but by the specific ISG program engaged, the timing of its induction, and the surrounding immune microenvironment. We propose that this conceptual framework may help reconcile apparently conflicting reports regarding the role of IFN-responsive neutrophils in cancer immunotherapy.

Our spatial transcriptomic analyses provide additional mechanistic insight into how neutrophil interferon programs may influence therapeutic outcome. Responding tumors exhibited coordinated enrichment of B cells, T cells, and dendritic cells consistent with TLSs, whereas IFN^hi^ neutrophils were markedly reduced and spatially excluded from these lymphocyte-rich immune niches. TLSs have emerged as important predictors of favorable responses to ICB because they support local antigen presentation, T-cell priming, and adaptive immune activation.^28,29,47,48^ Our findings raise the possibility that persistent IFN-responsive neutrophils interfere with the establishment or function of these specialized immune microenvironments. Whether these neutrophils actively disrupt TLS formation, suppress lymphocyte recruitment, or are excluded as a consequence of productive immune activation remains to be determined. Nevertheless, the reciprocal spatial relationship between IFNhi neutrophils and TLS-enriched regions suggests a previously unappreciated mechanism through which neutrophil interferon programs may restrain effective anti-tumor immunity.

Our genetic studies further establish that neutrophil-intrinsic interferon signaling contributes directly to immunotherapy resistance. Deletion of either IFNAR1 or IFNGR1 alone was insufficient to improve anti-PD1 efficacy, whereas simultaneous disruption of both receptors markedly enhanced therapeutic responses across multiple breast cancer models. This finding is consistent with the substantial overlap between type I and type II interferon signaling and their shared induction of STAT1-dependent ISG programs, allowing one pathway to partially compensate for loss of the other.^36,38^ Importantly, combined disruption also improved therapeutic efficacy in a neutrophil-enriched model exhibiting de novo resistance, suggesting that neutrophil interferon signaling contributes to both intrinsic and acquired resistance. These observations further support the development of strategies that selectively modulate pathogenic neutrophil interferon programs while preserving the essential anti-tumor functions of interferons in dendritic cells, T cells, and other immune populations.

Beyond regulating primary therapeutic response, our study identifies an unexpected role for neutrophil interferon signaling in controlling the durability of anti-tumor immunity. Successful anti-PD1 therapy generated robust T cell-dependent immunological memory capable of protecting mice from tumor rechallenge.^1,41,49^ However, neutrophil-enriched tumors impaired this established immune memory, whereas neutrophil depletion or disruption of neutrophil interferon signaling restored protective immunity. These findings extend the function of neutrophils beyond mediating primary resistance and suggest that persistent interferon-responsive neutrophils may also compromise the establishment and maintenance of long-term anti-tumor immune surveillance. Such effects may have important implications for preventing tumor recurrence and metastatic relapse following initially successful immunotherapy.

Several limitations warrant consideration. Although our findings were highly consistent across multiple preclinical breast cancer models, validation in patients receiving ICB will be essential to determine the clinical utility of circulating neutrophil ISG programs as predictive biomarkers. In addition, the downstream mechanisms through which IFN-responsive neutrophils suppress adaptive immunity remain incompletely understood. Potential mechanisms include PD-L1-mediated T-cell inhibition, altered chemokine production, disruption of TLS organization, or modulation of dendritic cell function. Finally, because interferon signaling performs indispensable functions in numerous immune cell populations, future therapeutic strategies should aim to selectively target pathogenic neutrophil interferon programs while preserving beneficial interferon responses elsewhere within the tumor microenvironment.

In summary, our study identifies an early neutrophil interferon program that precedes tumor response divergence, promotes resistance to PD-1 blockade, and limits durable anti-tumor immunity. More broadly, our findings suggest that the biological impact of neutrophil interferon signaling is determined not by interferon activation alone, but by the temporal dynamics and transcriptional composition of distinct interferon programs. This framework provides a unifying explanation for recent observations of both pro- and anti-tumor IFN-responsive neutrophil states and establishes a rationale for developing neutrophil-directed biomarkers and combination therapies to improve the depth and durability of cancer immunotherapy.

---

**Figure S1. Characterization of breast cancer models and early immune profiling supporting Figure 1**.

**(A, B)** Tumor growth curves of the neutrophil-enriched PyMT-N (A) and 2208L (B) breast cancer models following anti-PD1 or control IgG treatment. Both models exhibited limited responses to anti-PD1 therapy.

**(C)** Characteristics of the four preclinical breast cancer models used in this study, including molecular subtype (MES or NES), mouse strain, and genetic background.

**(D, E)** UMAP visualization (D) and quantification (E) of tumor-infiltrating immune cells from PyMT-N and PyMT-M tumors. PyMT-N tumor exhibited a neutrophil-enriched tumor microenvironment compared with PyMT-M tumor. N = 1 per group for PyMT-M and PyMT-N. **(F, G)** UMAP visualization (F) and quantification (G) of tumor-infiltrating immune cells from 2208L and T11 tumors. The 2208L model displayed increased neutrophil infiltration relative to the T11 model. N = 3 per group for T11 and 2208L.

**(H)** UMAP visualization of Day 5 peripheral blood immune cells from IgG-treated controls, anti-PD1 responders (PD1R), and anti-PD1 non-responders (PD1NR).

**(I)** Quantification of major immune cell populations in Day 5 peripheral blood. No significant differences in immune cell composition were observed between responders and non-responders at this time point. N = 3 per group for IgG-treated control, PD1R, and PD1NR.

**(J)** Dot plot showing expression of the neutrophil interferon-stimulated gene (ISG) signature identified in Figure 1 in Day 5 neutrophils. Sixteen ISGs were already downregulated in responders compared with non-responders, although only *Ifitm1* and *Ifitm3* reached statistical significance at this early time point.

**(K)** Dot plot illustrating expression of the Day 5 ISG signature across major peripheral blood immune cell populations. The identified ISGs were predominantly expressed in neutrophils, with *Ifitm1* exhibiting the greatest neutrophil specificity.

**Figure S2. Validation of spatial transcriptomic cell-type deconvolution using cell2location.**

**(A)** Quantification of B-cell, T-cell, and neutrophil abundance in Visium spatial transcriptomic datasets using the cell2location deconvolution algorithm. Consistent with the RCTD analyses presented in Figure 2, responder tumors (PD1R) exhibited significantly increased B-cell and T-cell abundance together with reduced neutrophil abundance compared with anti-PD1 non-responders (PD1NR) and IgG-treated controls. N = 2 per group for IgG-treated control, PD1R, and PD1NR tumors.

**(B)** Representative spatial distribution maps of B cells, T cells, and neutrophils generated using cell2location. Independent deconvolution confirmed enrichment of adaptive immune cells and reduced neutrophil abundance in responder tumors, validating the robustness of the spatial immune remodeling observed in Figure 2.

**Figure S3. Characterization and validation of IFN-responsive neutrophils in anti-PD1-responsive and -resistant tumors.**

**(A)** Representative spatial maps showing Ifitm1 expression inferred in individual cell types using cell2location deconvolution. Ifitm1 expression was predominantly localized to neutrophils, supporting its use as a representative marker of IFN-responsive neutrophils.

**(B)** Relative abundance of IFN^hi^ and IFN^low^ neutrophils estimated by cell2location in IgG-treated controls, anti-PD1 non-responders (PD1NR), and anti-PD1 responders (PD1R). Responding tumors exhibited significantly fewer IFN^hi^ neutrophils. N = 2 per group for IgG-treated control, PD1R, and PD1NR tumors.

**(C)** Pearson’s correlation analysis between IFN^hi^ or IFN^low^ neutrophils and B cells or T cells in IgG-treated controls, PD1NR, and PD1R tumors. IFN^hi^ neutrophils exhibited progressively reduced correlation with lymphocyte populations during successful anti-PD1 therapy, whereas IFN^low^ neutrophils showed stronger positive associations.

**(D)** Spearman’s correlation analysis between IFN^hi^ or IFN^low^ neutrophils and B cells or T cells in IgG-treated controls, PD1NR, and PD1R tumors, confirming the spatial relationships observed using Pearson’s correlation.

**(E)** Gene set enrichment analysis (GSEA) of neutrophils isolated from the anti-PD1-resistant 2208L and anti-PD1-responsive T11 tumor models. Hallmark pathways enriched in 2208L neutrophils included interferon-α response, interferon-γ response, etc.

**(F)** Comparison of the top enriched Hallmark pathways in tumor cells from the PyMT-N versus PyMT-M and 2208L versus T11 models. Interferon-α and interferon-γ response pathways were commonly enriched in both anti-PD1-resistant tumor models.

**(G)** Heatmap showing expression of interferon-stimulated genes (ISGs) in bone marrow neutrophils cocultured with PyMT-M or PyMT-N tumor cells. Coculture with PyMT-N cells induced expression of multiple ISGs, consistent with acquisition of an IFN-responsive neutrophil program.

**Figure S4. Validation of neutrophil-specific deletion of IFNAR1 and IFNGR1 in double knockout mice.**

**(A)** Flow cytometry gating strategy for analysis of IFNAR1 and IFNGR1 expression in immune cell populations isolated from tumors.

**(B)** Flow cytometry histograms showing IFNAR1 surface expression (mean fluorescence intensity, MFI) in neutrophils, Ly6C high monocytes, CD4⁺ T cells, and CD8⁺ T cells from IFNAR1F/F; IFNGR1F/F control mice and IFNAR1F/F; IFNGR1F/F; S100A8Cre mice. IFNAR1 expression was selectively reduced in neutrophils, whereas expression in monocytes and T-cell populations was preserved. N = 5 per group.

**(C)** Flow cytometry histograms showing IFNGR1 surface expression (MFI) in neutrophils, Ly6C high monocytes, CD4⁺ T cells, and CD8⁺ T cells from control and double knockout mice. IFNGR1 expression was specifically reduced in neutrophils but remained unchanged in the other immune cell populations. N = 5 per group.

**Figure S5. Successful immune checkpoint blockade generates durable systemic anti-tumor immunity.**

**(A)** Experimental design for evaluating systemic anti-tumor immunity following successful immune checkpoint blockade (ICB). Mice that completely rejected orthotopic E0771 mammary tumors after anti-PD1 treatment ("Cured") or age-matched untreated control mice ("Ctrl") received intracardiac injection of E0771 cells to induce experimental metastasis. Metastatic burden was assessed two weeks later.

**(B)** Representative images of adrenal and ovarian metastases following intracardiac injection of E0771 cells. Mice cured by anti-PD1 therapy didn’t develop metastatic lesions.

**(C)** Representative bioluminescence images showing metastatic tumor burden in control and cured mice at the indicated time points following intracardiac inoculation. Successful anti-PD1 therapy conferred durable systemic protection against metastatic tumor growth.

**(D)** Tumor growth following PyMT-M rechallenge in anti-PD1 responder (PD1R) mice treated with control IgG, T-cell depletion, B-cell depletion, or combined T- and B-cell depletion antibodies. Depletion of T cells, but not B cells, abrogated protective immune memory, indicating that durable anti-tumor immunity is primarily mediated by T cells.

**(E)** UMAP visualization of T-cell populations isolated from PyMT-N and PyMT-M tumors, showing the distribution of memory T cells and regulatory T cells (Tregs). PyMT-M tumors contained a greater proportion of memory T cells, whereas PyMT-N tumors were enriched for Tregs, consistent with impaired long-term anti-tumor immunity in the neutrophil-enriched tumor microenvironment.

## STAR METHODS

### Key resources table

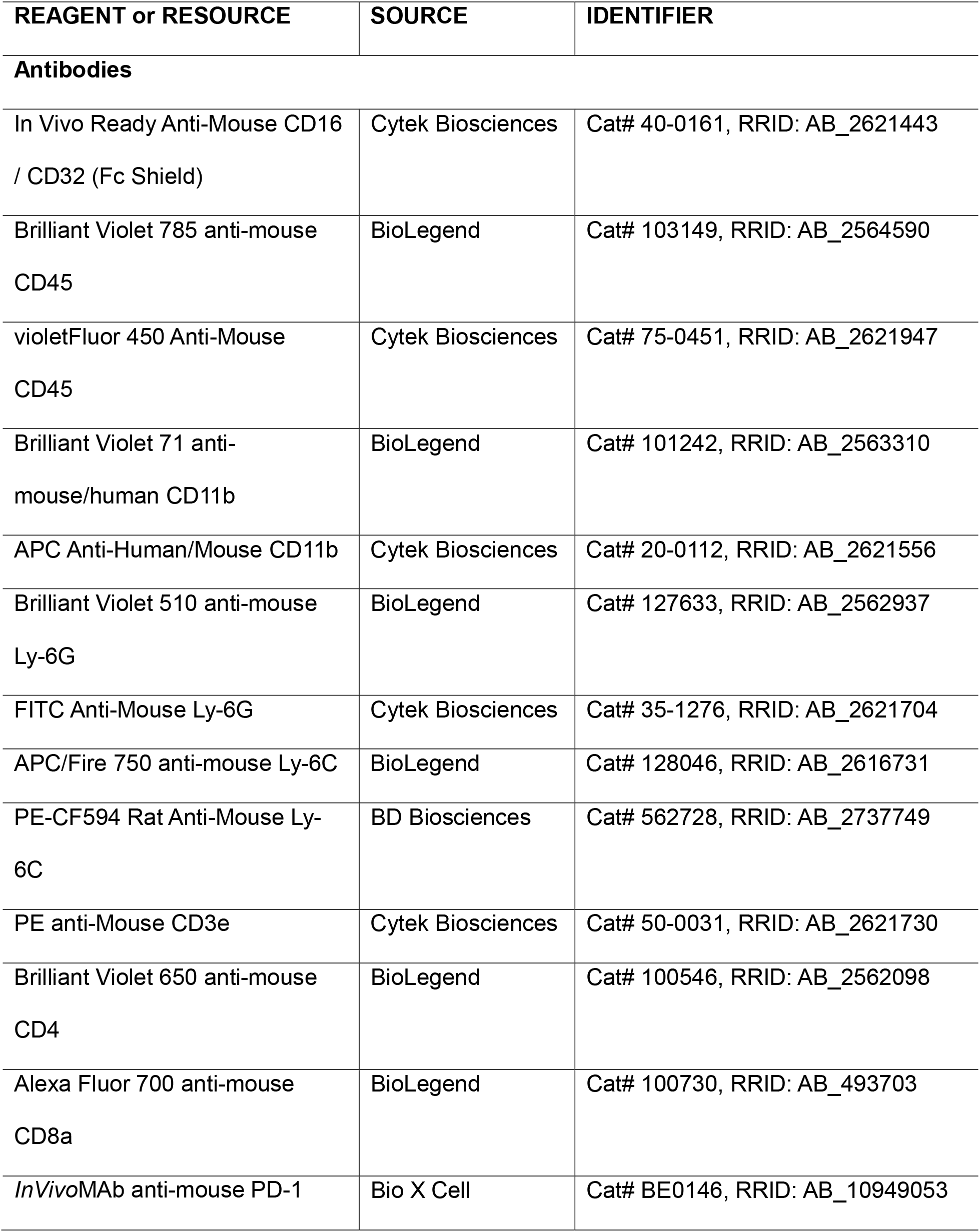
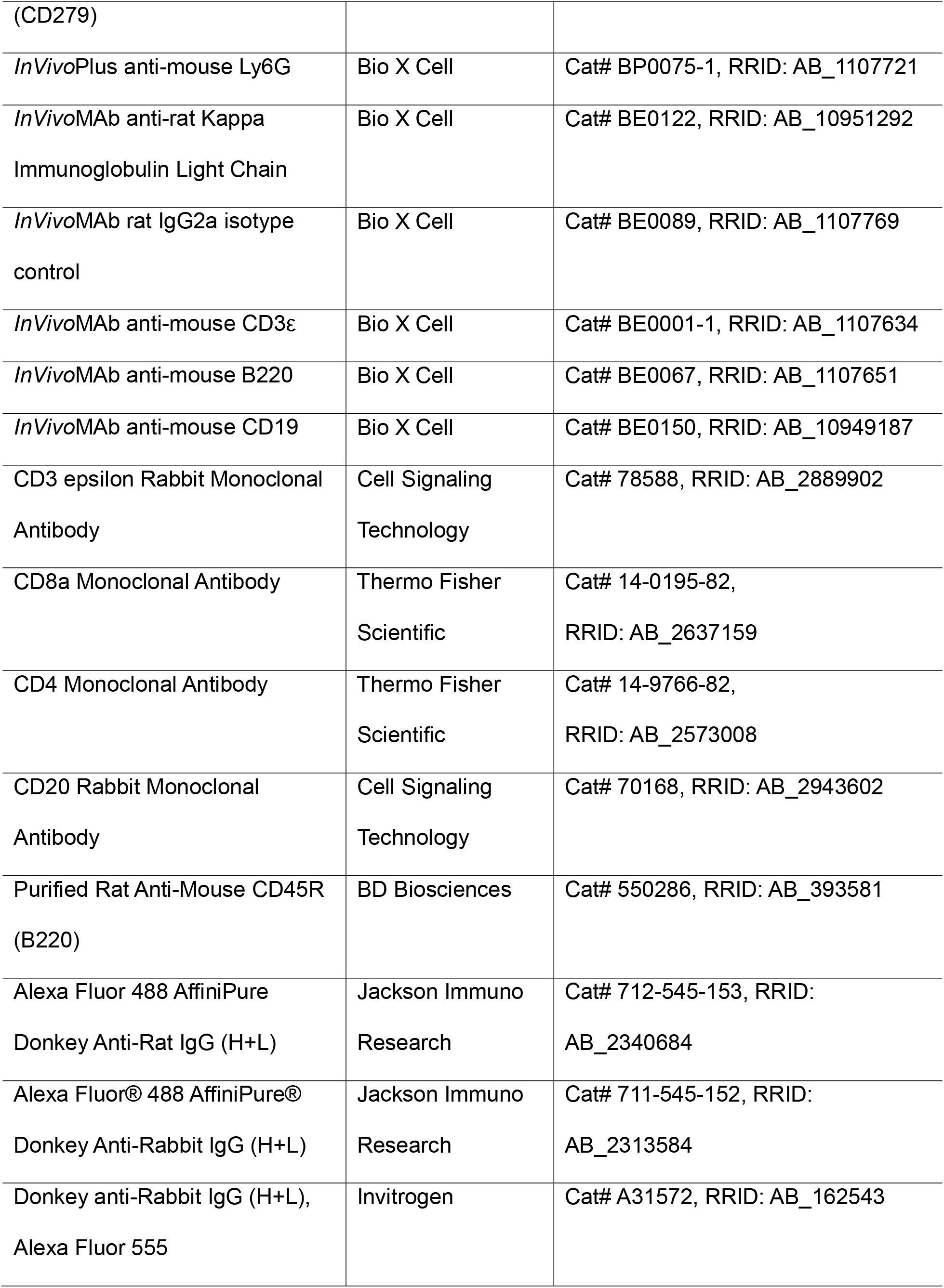
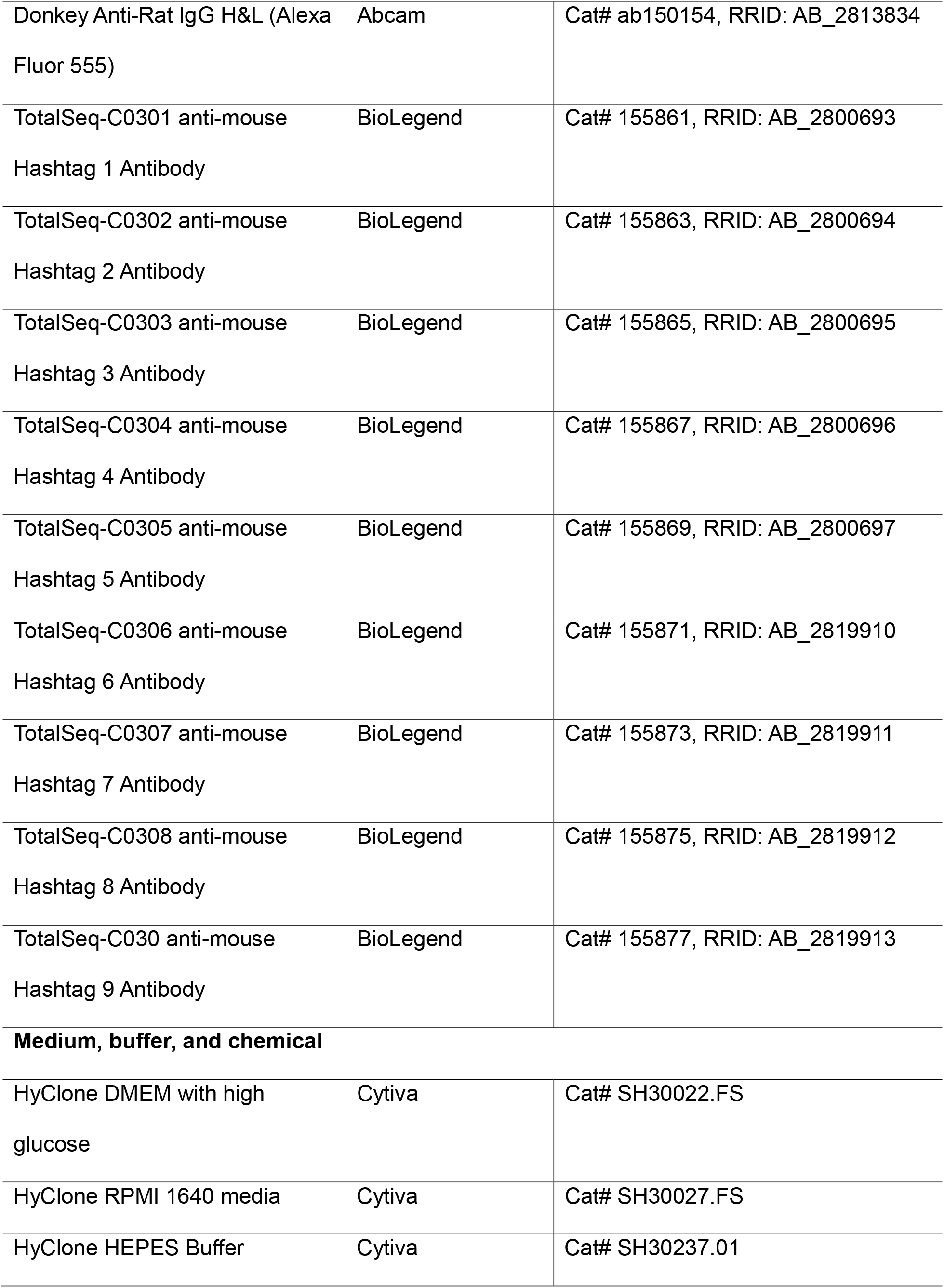
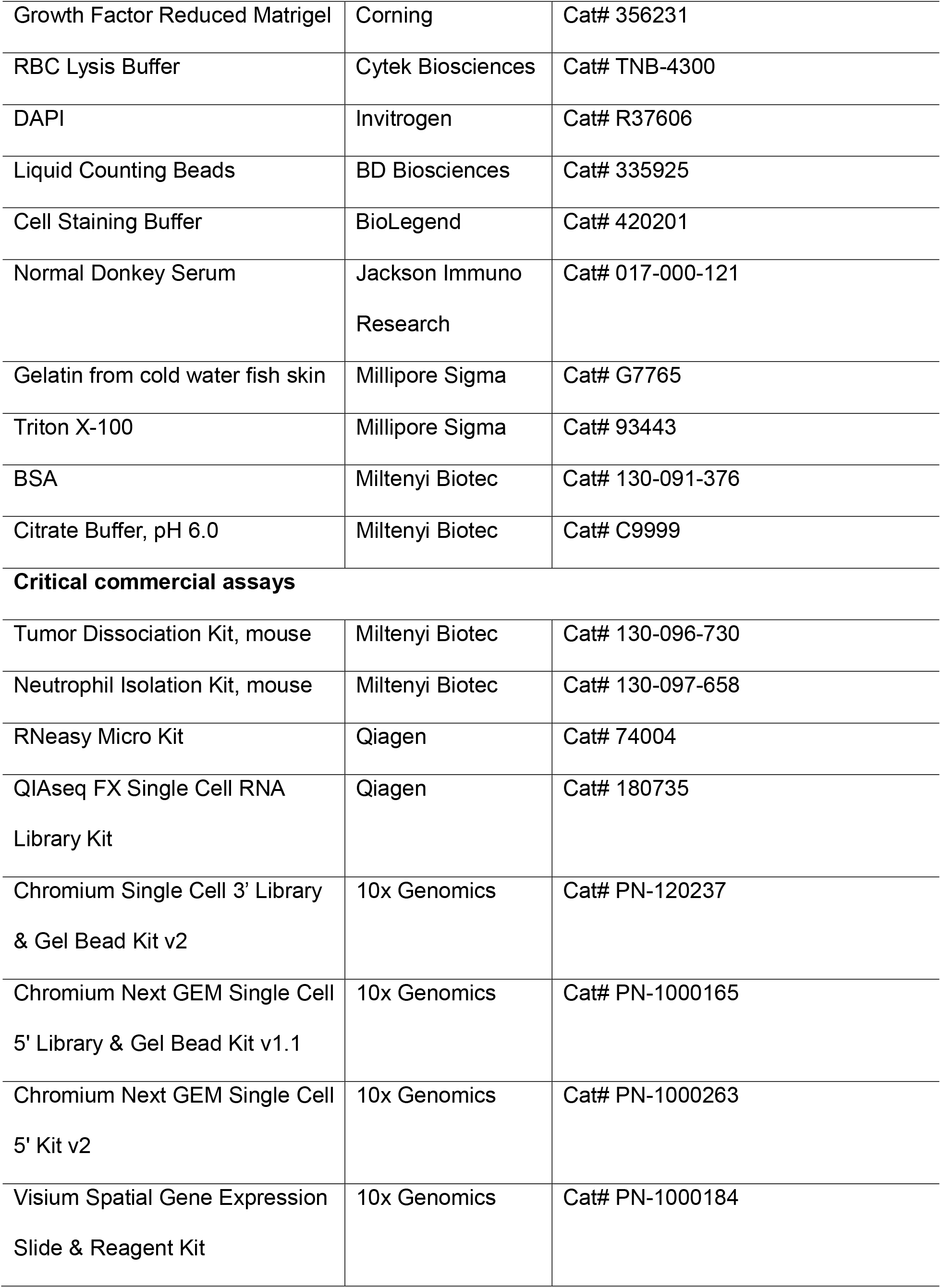
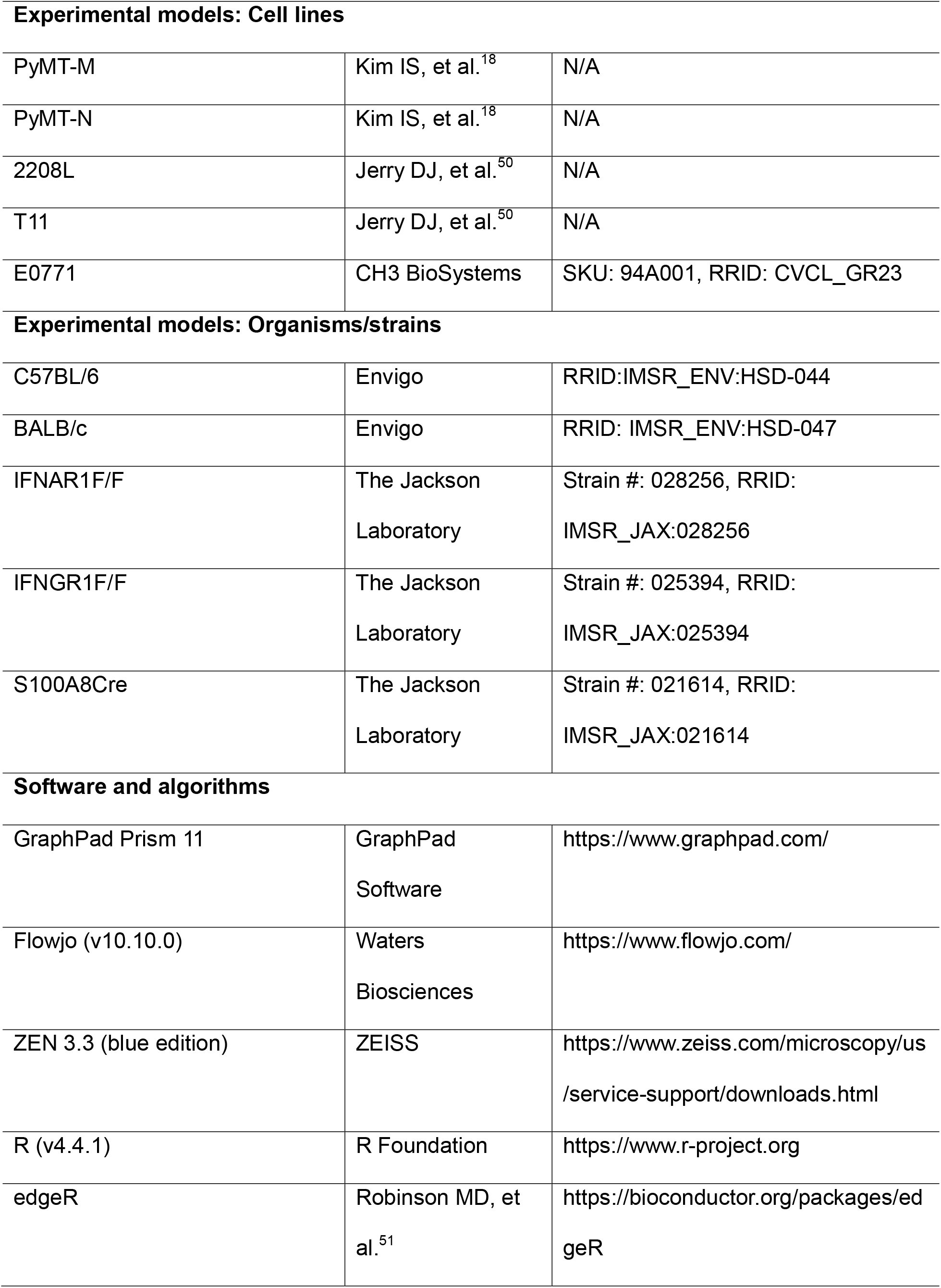
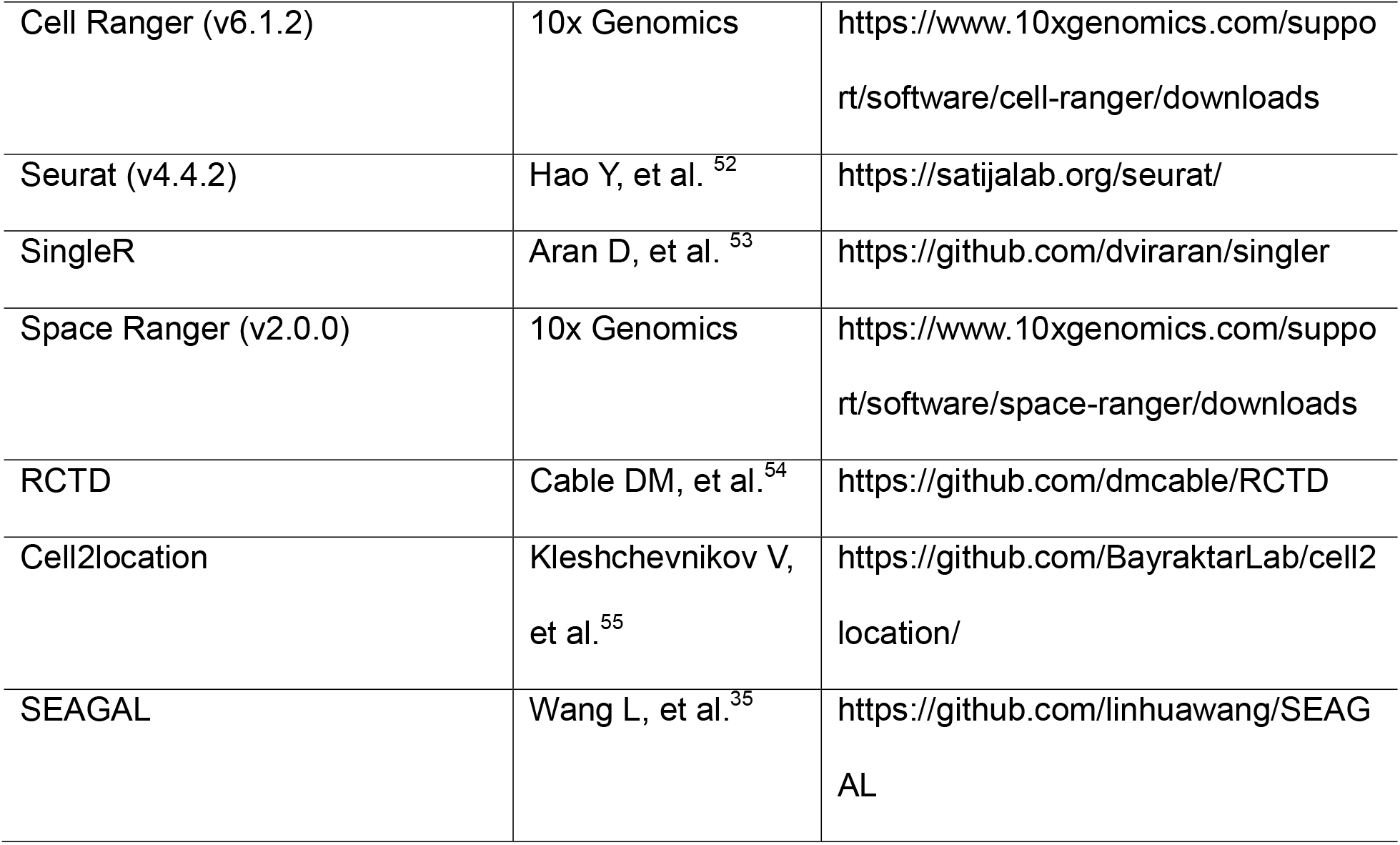

| REAGENT or RESOURCE | SOURCE | IDENTIFIER |
| --- | --- | --- |
| **Antibodies** |  |  |
| In Vivo Ready Anti-Mouse CD16 / CD32 (Fc Shield) | Cytek Biosciences | Cat# 40-0161, RRID: AB_2621443 |
| Brilliant Violet 785 anti-mouse CD45 | BioLegend | Cat# 103149, RRID: AB_2564590 |
| violetFluor 450 Anti-Mouse CD45 | Cytek Biosciences | Cat# 75-0451, RRID: AB_2621947 |
| Brilliant Violet 71 anti-mouse/human CD11b | BioLegend | Cat# 101242, RRID: AB_2563310 |
| APC Anti-Human/Mouse CD11b | Cytek Biosciences | Cat# 20-0112, RRID: AB_2621556 |
| Brilliant Violet 510 anti-mouse Ly-6G | BioLegend | Cat# 127633, RRID: AB_2562937 |
| FITC Anti-Mouse Ly-6G | Cytek Biosciences | Cat# 35-1276, RRID: AB_2621704 |
| APC/Fire 750 anti-mouse Ly-6C | BioLegend | Cat# 128046, RRID: AB_2616731 |
| PE-CF594 Rat Anti-Mouse Ly-6C | BD Biosciences | Cat# 562728, RRID: AB_2737749 |
| PE anti-Mouse CD3e | Cytek Biosciences | Cat# 50-0031, RRID: AB_2621730 |
| Brilliant Violet 650 anti-mouse CD4 | BioLegend | Cat# 100546, RRID: AB_2562098 |
| Alexa Fluor 700 anti-mouse CD8a | BioLegend | Cat# 100730, RRID: AB_493703 |
| *InVivo*MAb anti-mouse PD-1 | Bio X Cell | Cat# BE0146, RRID: AB_10949053 |
| (CD279) |  |  |
| *InVivo*Plus anti-mouse Ly6G | Bio X Cell | Cat# BP0075-1, RRID: AB_1107721 |
| *InVivo*MAb anti-rat Kappa<br>Immunoglobulin Light Chain | Bio X Cell | Cat# BE0122, RRID: AB_10951292 |
| *InVivo*MAb rat IgG2a isotype<br>control | Bio X Cell | Cat# BE0089, RRID: AB_1107769 |
| *InVivo*MAb anti-mouse CD3ε | Bio X Cell | Cat# BE0001-1, RRID: AB_1107634 |
| *InVivo*MAb anti-mouse B220 | Bio X Cell | Cat# BE0067, RRID: AB_1107651 |
| *InVivo*MAb anti-mouse CD19 | Bio X Cell | Cat# BE0150, RRID: AB_10949187 |
| CD3 epsilon Rabbit Monoclonal<br>Antibody | Cell Signaling<br>Technology | Cat# 78588, RRID: AB_2889902 |
| CD8a Monoclonal Antibody | Thermo Fisher<br>Scientific | Cat# 14-0195-82,<br>RRID: AB_2637159 |
| CD4 Monoclonal Antibody | Thermo Fisher<br>Scientific | Cat# 14-9766-82,<br>RRID: AB_2573008 |
| CD20 Rabbit Monoclonal<br>Antibody | Cell Signaling<br>Technology | Cat# 70168, RRID: AB_2943602 |
| Purified Rat Anti-Mouse CD45R<br>(B220) | BD Biosciences | Cat# 550286, RRID: AB_393581 |
| Alexa Fluor 488 AffiniPure<br>Donkey Anti-Rat IgG (H+L) | Jackson Immuno<br>Research | Cat# 712-545-153, RRID:<br>AB_2340684 |
| Alexa Fluor® 488 AffiniPure®<br>Donkey Anti-Rabbit IgG (H+L) | Jackson Immuno<br>Research | Cat# 711-545-152, RRID:<br>AB_2313584 |
| Donkey anti-Rabbit IgG (H+L),<br>Alexa Fluor 555 | Invitrogen | Cat# A31572, RRID: AB_162543 |
| Donkey Anti-Rat IgG H&L (Alexa Fluor 555) | Abcam | Cat# ab150154, RRID: AB_2813834 |
| TotalSeq-C0301 anti-mouse Hashtag 1 Antibody | BioLegend | Cat# 155861, RRID: AB_2800693 |
| TotalSeq-C0302 anti-mouse Hashtag 2 Antibody | BioLegend | Cat# 155863, RRID: AB_2800694 |
| TotalSeq-C0303 anti-mouse Hashtag 3 Antibody | BioLegend | Cat# 155865, RRID: AB_2800695 |
| TotalSeq-C0304 anti-mouse Hashtag 4 Antibody | BioLegend | Cat# 155867, RRID: AB_2800696 |
| TotalSeq-C0305 anti-mouse Hashtag 5 Antibody | BioLegend | Cat# 155869, RRID: AB_2800697 |
| TotalSeq-C0306 anti-mouse Hashtag 6 Antibody | BioLegend | Cat# 155871, RRID: AB_2819910 |
| TotalSeq-C0307 anti-mouse Hashtag 7 Antibody | BioLegend | Cat# 155873, RRID: AB_2819911 |
| TotalSeq-C0308 anti-mouse Hashtag 8 Antibody | BioLegend | Cat# 155875, RRID: AB_2819912 |
| TotalSeq-C030 anti-mouse Hashtag 9 Antibody | BioLegend | Cat# 155877, RRID: AB_2819913 |
| **Medium, buffer, and chemical** |  |  |
| HyClone DMEM with high glucose | Cytiva | Cat# SH30022.FS |
| HyClone RPMI 1640 media | Cytiva | Cat# SH30027.FS |
| HyClone HEPES Buffer | Cytiva | Cat# SH30237.01 |
| Growth Factor Reduced Matrigel | Corning | Cat# 356231 |
| RBC Lysis Buffer | Cytek Biosciences | Cat# TNB-4300 |
| DAPI | Invitrogen | Cat# R37606 |
| Liquid Counting Beads | BD Biosciences | Cat# 335925 |
| Cell Staining Buffer | BioLegend | Cat# 420201 |
| Normal Donkey Serum | Jackson Immuno<br>Research | Cat# 017-000-121 |
| Gelatin from cold water fish skin | Millipore Sigma | Cat# G7765 |
| Triton X-100 | Millipore Sigma | Cat# 93443 |
| BSA | Miltenyi Biotec | Cat# 130-091-376 |
| Citrate Buffer, pH 6.0 | Miltenyi Biotec | Cat# C9999 |
| **Critical commercial assays** |  |  |
| Tumor Dissociation Kit, mouse | Miltenyi Biotec | Cat# 130-096-730 |
| Neutrophil Isolation Kit, mouse | Miltenyi Biotec | Cat# 130-097-658 |
| RNeasy Micro Kit | Qiagen | Cat# 74004 |
| QIAseq FX Single Cell RNA<br>Library Kit | Qiagen | Cat# 180735 |
| Chromium Single Cell 3' Library<br>& Gel Bead Kit v2 | 10x Genomics | Cat# PN-120237 |
| Chromium Next GEM Single Cell<br>5' Library & Gel Bead Kit v1.1 | 10x Genomics | Cat# PN-1000165 |
| Chromium Next GEM Single Cell<br>5' Kit v2 | 10x Genomics | Cat# PN-1000263 |
| Visium Spatial Gene Expression<br>Slide & Reagent Kit | 10x Genomics | Cat# PN-1000184 |

| **Experimental models: Cell lines** |  |  |
| --- | --- | --- |
| PyMT-M | Kim IS, et al.<sup>18</sup> | N/A |
| PyMT-N | Kim IS, et al.<sup>18</sup> | N/A |
| 2208L | Jerry DJ, et al.<sup>50</sup> | N/A |
| T11 | Jerry DJ, et al.<sup>50</sup> | N/A |
| E0771 | CH3 BioSystems | SKU: 94A001, RRID: CVCL_GR23 |
| **Experimental models: Organisms/strains** |  |  |
| C57BL/6 | Envigo | RRID:IMSR_ENV:HSD-044 |
| BALB/c | Envigo | RRID: IMSR_ENV:HSD-047 |
| IFNAR1F/F | The Jackson Laboratory | Strain #: 028256, RRID: IMSR_JAX:028256 |
| IFNGR1F/F | The Jackson Laboratory | Strain #: 025394, RRID: IMSR_JAX:025394 |
| S100A8Cre | The Jackson Laboratory | Strain #: 021614, RRID: IMSR_JAX:021614 |
| **Software and algorithms** |  |  |
| GraphPad Prism 11 | GraphPad Software | https://www.graphpad.com/ |
| Flowjo (v10.10.0) | Waters Biosciences | https://www.flowjo.com/ |
| ZEN 3.3 (blue edition) | ZEISS | https://www.zeiss.com/microscopy/us/service-support/downloads.html |
| R (v4.4.1) | R Foundation | https://www.r-project.org |
| edgeR | Robinson MD, et al.<sup>51</sup> | https://bioconductor.org/packages/edgeR |
| Cell Ranger (v6.1.2) | 10x Genomics | https://www.10xgenomics.com/support/software/cell-ranger/downloads |
| Seurat (v4.4.2) | Hao Y, et al.<sup>52</sup> | https://satijalab.org/seurat/ |
| SingleR | Aran D, et al.<sup>53</sup> | https://github.com/dviraran/singler |
| Space Ranger (v2.0.0) | 10x Genomics | https://www.10xgenomics.com/support/software/space-ranger/downloads |
| RCTD | Cable DM, et al.<sup>54</sup> | https://github.com/dmcable/RCTD |
| Cell2location | Kleshchevnikov V, et al.<sup>55</sup> | https://github.com/BayraktarLab/cell2location/ |
| SEAGAL | Wang L, et al.<sup>35</sup> | https://github.com/linhuawang/SEAGAL |

## EXPERIMENTAL MODEL AND STUDY PARTICIPANT DETAILS

### Mice

All animal experiments were conducted in accordance with protocols approved by the Institutional Animal Care and Use Committee of Baylor College of Medicine (Houston, TX) and in compliance with all relevant ethical regulations regarding animal research. Wild-type BALB/c and C57/B6 mice were purchased from Envigo. IFNAR1F/F (Strain #: 028256), IFNGR1F/F (Strain #: 025394), and S100A8Cre (Strain #: 021614) mice were purchased from The Jackson Laboratory. IFNAR1F/F; IFNGR1F/F mice were generated by breeding IFNAR1F/F with IFNGR1F/F mice. All mice were kept on a 12 h light–dark cycle at the room temperature was maintained between 20–24 °C, and the humidity was maintained at 40–70%.

### Tumor cell lines

Murine TNBC lines PyMT-M (C57BL/6), PyMT-N (C57BL/6) 2208L (BALB/c), and T11 (BALB/c) were cultured in DMEM/high glucose medium (Cytivia) containing 10% FBS (Gibco) and antibiotics. E0771 (C57BL/6) cells were cultured in RPMI-1640 medium (Cytivia) containing 10% FBS and 1× HEPES (Cytivia). All cell lines were cultured in a 5% CO_2_ 37°C incubator. All cell lines were routinely tested for *Mycoplasma* contamination and confirmed to be negative before use.

## METHOD DETAILS

### Orthotopic tumor implantation and rechallenge

Female mice, 6–8 weeks of age, were used for orthotopic tumor implantation. Tumor cells were harvested using 0.25% trypsin (Corning), washed with PBS (Lonza), counted, and resuspended in PBS. PyMT-M, PyMT-N, E0771, 2208L, or T11 cells (2 × 10^5^ cells in 20 μL PBS) were mixed 1:1 with growth factor-reduced Matrigel (Corning) and injected into the right fourth mammary fat pad as previously described.^18^ For bilateral tumor models, 2 × 10^5^ tumor cells were implanted into each of the left and right fourth mammary fat pads.

For tumor rechallenge experiments, 2 × 10^5^ tumor cells were implanted into the contralateral (left) fourth mammary fat pad to assess systemic anti-tumor immunological memory.

### Intracardiac Injection

For intracardiac injection, tumor cells (1 × 10^5^ cells in 100 μL PBS) were injected into the left ventricle of anesthetized mice using a 26-gauge needle (BD Biosciences), as previously described.^56^

### In vivo drug treatment and immune cell depletion

Tumor cells were orthotopically implanted on day 0, and mice were randomized to treatment groups after tumor implantation. Anti-PD-1 antibody (200 μg per mouse; Bio X Cell) was administered intraperitoneally beginning on day 3 and every 3 days thereafter for a total of six doses.^57^ Rat IgG2α isotype antibody (Bio X Cell) was used as the control.

For neutrophil depletion, mice received anti-Ly6G antibody (25 μg per mouse; Bio X Cell) daily from days 0–18. Anti-kappa light chain antibody (50 μg per mouse; Bio X Cell) was injected every other day over the same period.^58,59^

For T cell depletion, anti-CD3ε antibody (200 μg per mouse; Bio X Cell) was administered beginning 4 days before tumor rechallenge through day 2 after rechallenge, followed by treatment every 3 days from days 5–18.^60^

For B cell depletion, mice received a combination of anti-B220 (75 μg per mouse; Bio X Cell) and anti-CD19 (75 μg per mouse; Bio X Cell) antibodies daily beginning 5 days before tumor rechallenge through day 2 after rechallenge, followed by treatment every 3 days from days 5–18. In the meantime, anti-kappa light chain antibody (100 μg per mouse; Bio X Cell) was administered every other day beginning 4 days before rechallenge through day 2, followed by treatment every 3 days from days 5–18.^61^ All antibodies used for immune cell depletion were administered intraperitoneally.

### Tissue collection and dissociation

Peripheral blood was collected using microhematocrit capillary tubes (Kimble). Red blood cells (RBC) were lysed twice with RBC lysis buffer for 5 minutes on ice each time.

For isolation of splenic immune cells, spleens were mechanically dissociated and passed through a 70-μm cell strainer. Red blood cells were lysed with RBC lysis buffer for 5 minutes on ice.

For tumor tissues, tumors were collected in ice-cold PBS and dissociated using the Mouse Tumor Dissociation Kit (Miltenyi Biotec) and gentleMACS Dissociator (Miltenyi Biotec) according to the manufacturer’s instructions. Following dissociation, samples were passed through a 70-μm cell strainer to obtain single-cell suspensions. Red blood cells were lysed with RBC lysis buffer for 1 minute on ice.

### Immunofluorescence staining and imaging

Formalin-fixed, paraffin-embedded tissues were prepared and sectioned by the Breast Center Pathology Core at Baylor College of Medicine. Tissue sections were deparaffinized in xylene and rehydrated through a graded ethanol series. Heat-induced antigen retrieval was performed by boiling tissue sections in 10 mM sodium citrate buffer (pH 6.0) (Millipore Sigma) for 20 min. Sections were subsequently blocked and permeabilized in PBS containing 10% normal donkey serum (Jackson ImmunoResearch), 2% fish gelatin (Millipore Sigma), and 0.1% Triton X-100 (Millipore Sigma).

Sections were incubated with primary antibodies overnight at 4°C. After washing with PBS, sections were incubated with the appropriate fluorophore-conjugated secondary antibodies for 2 h at room temperature. Antibodies used in this study are listed in the Key Resources Table.

Fluorescence images were acquired using a ZEISS LSM 780 or LSM 880 confocal microscope with 16× line averaging as describe before.^62^ Images were processed and analyzed using ZEN 3.3 (blue edition) software (ZEISS).

### Flow cytometry

Single-cell suspensions were incubated with Fc receptor blocking antibody (Cytek Biosciences) for 10 minutes on ice. Cells were subsequently stained with fluorophore-conjugated antibodies in FACS buffer (PBS containing 2% FBS) for 25 minutes on ice in the dark. After washing, cells were resuspended in FACS buffer containing DAPI (BD Biosciences) for exclusion of dead cells. Data were acquired on a BD LSRFortessa or LSR II flow cytometer (BD Biosciences), and analyzed using FlowJo v10.10.0. Antibodies used for flow cytometry are listed in the Key Resources Table.

### Neutrophil isolation, coculture, and RNA sequencing

Neutrophils were isolated from bone marrow (BM) by negative selection using the mouse Neutrophil Isolation Kit (Miltenyi Biotec). BM cells were collected from the femurs and tibias and subjected to RBC lysis. Cells were labeled with the neutrophil enrichment cocktail and streptavidin-conjugated magnetic beads according to the manufacturer’s instructions. Labeled cells were passed through LS columns (Miltenyi Biotec), and negatively selected neutrophils were collected from the flow-through.

For coculture experiments, BM-derived neutrophils were cocultured with tumor cells at a neutrophil-to-tumor cell ratio of 5:1 in DMEM containing 0.5% FBS.^63^ Following overnight coculture, cells were collected and stained with antibodies against CD45, CD11b, Ly6G, and Ly6C to distinguish neutrophils from tumor cells. DAPI (Invitrogen) was used to exclude dead cells. Counting beads (BD Biosciences) were included for absolute cell quantification by flow cytometry. For downstream RNA sequencing, neutrophils (DAPI^−^CD45^+^CD11b^+^Ly6G^+^Ly6C^med-low^) were purified by fluorescence-activated cell sorting (FACS) using a FACSAria II Cell Sorter (BD Biosciences). Antibodies and reagents used in these experiments are listed in the Key Resources Table.

For RNA sequencing, total RNA was extracted from FACS-purified neutrophils using the RNeasy Micro Kit (QIAGEN). RNA-seq libraries were generated from 150 pg total RNA using the QIAseq FX Single Cell RNA Library Kit (QIAGEN) according to the manufacturer’s instructions. Raw sequencing reads were filtered to remove reads containing adapter contamination, >10% undetermined nucleotides, or >50% low-quality nucleotides. Read quality was assessed using FastQC before and after preprocessing. Filtered reads were aligned to the mouse reference genome (GRCm38/mm10), and gene-level expression was quantified to generate a count matrix for downstream analysis in R. Differential expression between neutrophils cocultured with PyMT-N and PyMT-M cells was assessed using the edgeR package.

### Bioluminescence imaging

For *in vivo* bioluminescence imaging, mice were anesthetized and administered 100 μL D-luciferin (15 mg/mL; GoldBio, cat. #LUCK-100) by retro-orbital injection. Mice were imaged immediately after D-luciferin administration using an IVIS Lumina imaging system (Revvity). Imaging was performed every 3 days.

### Library preparation and cell hashing for scRNA-seq

Tumor and peripheral blood samples were processed into single-cell suspensions as described in the “Tissue collection and dissociation” section.

For tumor-infiltrating immune cell scRNA-seq, DAPI^−^CD45^+^ were sorted and immediately submitted to the Single Cell Genomics Core at Baylor College of Medicine for library preparation. Single-cell gene expression libraries from T11 and 2208L tumor immune cells were generated using the Chromium Single Cell 3’ Library & Gel Bead Kit v2 (10x Genomics), whereas libraries from PyMT-M and PyMT-N tumors immune cells were generated using the Chromium Next GEM Single Cell 5′ v1.1 platform (10x Genomics).

For peripheral blood scRNA-seq, following RBC lysis, cells were counted and samples from individual mice were labeled with TotalSeq-C hashtag antibodies (BioLegend) according to the manufacturer’s instructions. Hashtag-labeled samples were pooled and single-cell gene expression libraries were generated using the Chromium Next GEM Single Cell 5′ v2 platform (10x Genomics).

All libraries were prepared by the Single Cell Genomics Core at Baylor College of Medicine according to the manufacturer’s instructions. Following library quality control, libraries were sequenced on a NovaSeq 6000 system (Illumina).

### Pre-processing and analysis of scRNA-seq data

Raw sequencing data were processed using Cell Ranger (10x Genomics) to generate gene expression and feature-barcode count matrices. Subsequent analyses were performed in R using Seurat.

For peripheral blood scRNA-seq, gene expression and hashtag oligonucleotide (HTO) count matrices were imported into Seurat. HTO counts were normalized using centered log-ratio transformation, and individual samples were demultiplexed using the HTODemux function with a positive quantile of 0.99. Cells classified as singlets were retained for downstream analysis. Quality control was performed by retaining cells with >200 and <3,000 detected genes and <5% mitochondrial transcripts. Mitochondrial genes (*mt-*), ribosomal protein genes (*Rpl* and *Rps*), and predicted genes beginning with *Gm* were excluded from downstream analysis. Gene expression data were processed using the standard Seurat workflow, including normalization, identification of variable features, scaling, PCA, graph-based clustering, and UMAP visualization.

For tumor-infiltrating immune-cell scRNA-seq, quality control was performed using the same criteria described above. T11 and 2208L datasets and PyMT-M and PyMT-N datasets were analyzed separately. For each pair, data were normalized, variable features were identified, and the data were scaled and subjected to PCA before integration in Seurat. Following integration, graph-based clustering was performed, and cell populations were visualized using UMAP.

For all scRNA-seq datasets, cell clusters were annotated using SingleR ^53^ with the ImmGen reference dataset ^64^ together with the expression of canonical lineage markers. Related clusters were grouped into major immune-cell populations for downstream analyses. Cell-type frequencies were calculated for each individual sample as the proportion of cells assigned to each annotated population. For peripheral blood samples, cells were assigned to control, anti-PD-1 responder, or anti-PD-1 non-responder groups based on sample-specific hashtag identities and treatment responses. For tumor datasets, immune-cell compositions were compared between T11 and 2208L tumors and between PyMT-M and PyMT-N tumors. Differential expression analyses between the indicated experimental groups were performed within specific cell populations using the FindMarkers function in Seurat. Gene set enrichment analysis (GSEA) was subsequently performed using ranked gene lists derived from differential expression analyses and Hallmark gene sets from the Molecular Signatures Database (MSigDB). Normalized enrichment scores (NES), *P* values, and adjusted *P* values were used to assess pathway enrichment.

### Spatial transcriptomics library preparation

PyMT-M tumors were collected from the left mammary fat pad of mice bearing bilateral tumors and treated with IgG control or anti-PD-1 antibody at day 8. Tumors were frozen and cryosectioned at 10 µm thickness. Tissue sections were placed onto Visium Spatial Gene Expression slides, and tissue permeabilization, spatially barcoded cDNA synthesis, cDNA amplification, and library preparation were performed using the Visium Spatial Gene Expression Slide & Reagent Kit (10x Genomics) according to the manufacturer’s instructions. Following library quality control, libraries were sequenced and raw sequencing data were processed using Space Ranger (10x Genomics) to generate spatially resolved gene-expression count matrices for downstream analysis.

### Spatial transcriptomic cell-type deconvolution by RCTD

Cell-type composition of Visium (10x Genomics) spatial transcriptomic data was estimated using Robust Cell Type Decomposition (RCTD) implemented in the spacexr package.^54^ A single-cell reference was generated by integrating PyMT-M tumor-infiltrating and peripheral blood immune-cell scRNA-seq datasets. Cell types in the integrated reference were annotated using SingleR with the ImmGen reference dataset and canonical lineage markers, as described above. Raw RNA counts and corresponding cell-type annotations from the single-cell reference were used to construct the RCTD reference.

For each spatial transcriptomic sample, raw gene-expression counts, total UMI counts, and spatial coordinates for individual spots were used as input. Genes shared between the spatial transcriptomic data and the single-cell reference were retained for deconvolution. RCTD was performed independently for each tissue section using doublet_mode = "full" and a minimum cell-type representation of five cells in the single-cell reference. The resulting cell-type weights at individual spatial spots were used to estimate cell-type composition for downstream comparisons and spatial visualization.

Differentially expressed genes were identified from pairwise comparisons among anti-PD-1 responder, anti-PD-1 non-responder, and IgG-treated control tumors. and were subjected to Gene Ontology (GO) enrichment analysis using the Biological Process ontology to identify associated biological processes.

To specifically examine the spatial distribution of IFN-associated neutrophils, an IFN-response score was calculated for each neutrophil in the single-cell reference using interferon-stimulated genes identified from the day 8 PyMT-M peripheral blood scRNA-seq analysis. Expression of individual genes was scaled across neutrophils and combined to generate an aggregate IFN-response score. Neutrophils were subsequently classified into IFN-high and IFN-low populations using the 25th percentile of the IFN-response score as a threshold. These annotations were incorporated into the single-cell reference, and RCTD was repeated to estimate the spatial distribution of IFN-high and IFN-low neutrophils.

### Spatial transcriptomic deconvolution by cell2location

As an independent approach to validate spatial cell-type composition, Visium spatial transcriptomic data were analyzed using cell2location.^55^ The same annotated scRNA-seq reference used for RCTD was used to derive cell-type expression signatures. Genes were filtered based on their expression across reference cells, and cell-type expression signatures were inferred using a negative binomial regression model, with cell-type annotation and sample batch information incorporated into the model. Posterior mean expression signatures for each cell population were used for subsequent spatial mapping.

For spatial mapping, raw Visium gene-expression counts were used as input. Mitochondrial genes were excluded, and the spatial transcriptomic data and reference expression signatures were restricted to shared genes. Individual spatial samples were included as a batch covariate. The cell2location spatial model was fitted with an expected abundance of five cells per spatial location (N_cells_per_location = 5) and detection_alpha = 20. Posterior cell-type abundance estimates were generated from 1,000 posterior samples.

The 5th percentile of the posterior abundance distribution (q05_cell_abundance_w_sf) was used as a conservative estimate of cell-type abundance at each spatial location. Relative abundance was calculated by normalizing the estimated abundance of each cell type to the summed abundance of all inferred cell populations within each spot. These estimates were used for comparisons among experimental groups and for visualization of cell-type distributions across the corresponding tissue sections.

### Spatial correlation analysis

Spatial associations among immune-cell populations were assessed using pairwise correlation analyses across Visium spots. For each cell type, expression of the corresponding marker-gene signature was summarized at individual spatial locations, and pairwise Spearman’s correlation coefficients were calculated within each tissue section. Correlation matrices were generated for major immune-cell populations and compared across experimental groups.

To quantify spatial associations involving IFN-associated neutrophils, Spatial Enrichment Analysis of Gene Associations using *L*-index (SEAGAL)^35^ was applied to the Visium spatial transcriptomic data. Cell-type marker-gene signatures were used to derive spatial scores for IFN-high neutrophils, IFN-low neutrophils, B cells, and T cells, and pairwise spatial associations were quantified using the *L*-index for each tissue section.

As independent validation, Pearson’s and Spearman’s correlation coefficients were calculated between the spatial marker-signature scores of IFN-high or IFN-low neutrophils and those of B or T cells across Visium spots within each tissue section. Correlation analyses were performed separately for individual samples and summarized across experimental groups.

### Quantification and statistical analysis

For *in vivo* experiments, individual mice were considered biological replicates, and sample sizes are indicated in the figures or figure legends. No statistical method was used to predetermine sample sizes, and all biologically independent samples were included in the analyses. Data were analyzed using GraphPad Prism. Tumor growth curves were analyzed using two-way ANOVA or mixed-effects analysis, as appropriate. Survival was analyzed using Kaplan–Meier analysis with the log-rank (Mantel–Cox) test. Additional statistical tests, where applicable, are specified in the corresponding figure legends.

## Acknowledgements

X. H.-F. Zhang is supported by US Department of Defense DAMD HT9425-23-1-0493, NCI R01CA183878, NCI R01CA251950, NCI R01CA221946, NCI R01CA227904, NCI U01CA253533, NCI P50CA186784, and Breast Cancer Research Foundation. Y. Gao is supported by NCI R00CA279899. The sequencing performed at the Single Cell Genomics Core and Genomic and RNA Profiling Core at BCM partially was supported by NIH (S10OD023469, S10OD025240, S10-OD032185), P30EY002520 and CPRIT RP200504. The flow cytometry analysis was supported by the Cytometry and Cell Sorting Core at Baylor College of Medicine with funding from the CPRIT Core Facility Support Award (CPRIT-RP180672), the NIH (CA125123 and RR024574) and the assistance of Joel M. Sederstrom. The confocal imaging was supported by the Optical Imaging & Vital Microscopy (OiVM) core at BCM.

## Author contributions

X.H.-F.Z., Y.G., and E.A. developed the concept, designed experiments, and wrote the manuscript. Y.G., E.A., L.W., Z.X., performed and analyzed the animal studies and *in vitro* experiments. K.L. and L.S.E. conducted tissue lipid characterization. Y.S. and F.L. performed scRNA-seq and bioinformatic analysis. B.W. and P.L.L. performed and analyzed the lipidomics experiments. J.L. and D.G.E. contributed to maintaining mouse strains. L.W., Z.X., and D.G.E assisted in animal work. Other authors contributed to manuscript editing. X.H.-F.Z. supervised the research.

