## Supplementary figures and images for "Early dynamics of a neutrophil interferon program drive immunotherapy resistance and limit durable anti-tumor immunity"

### Supplemental Fig 1

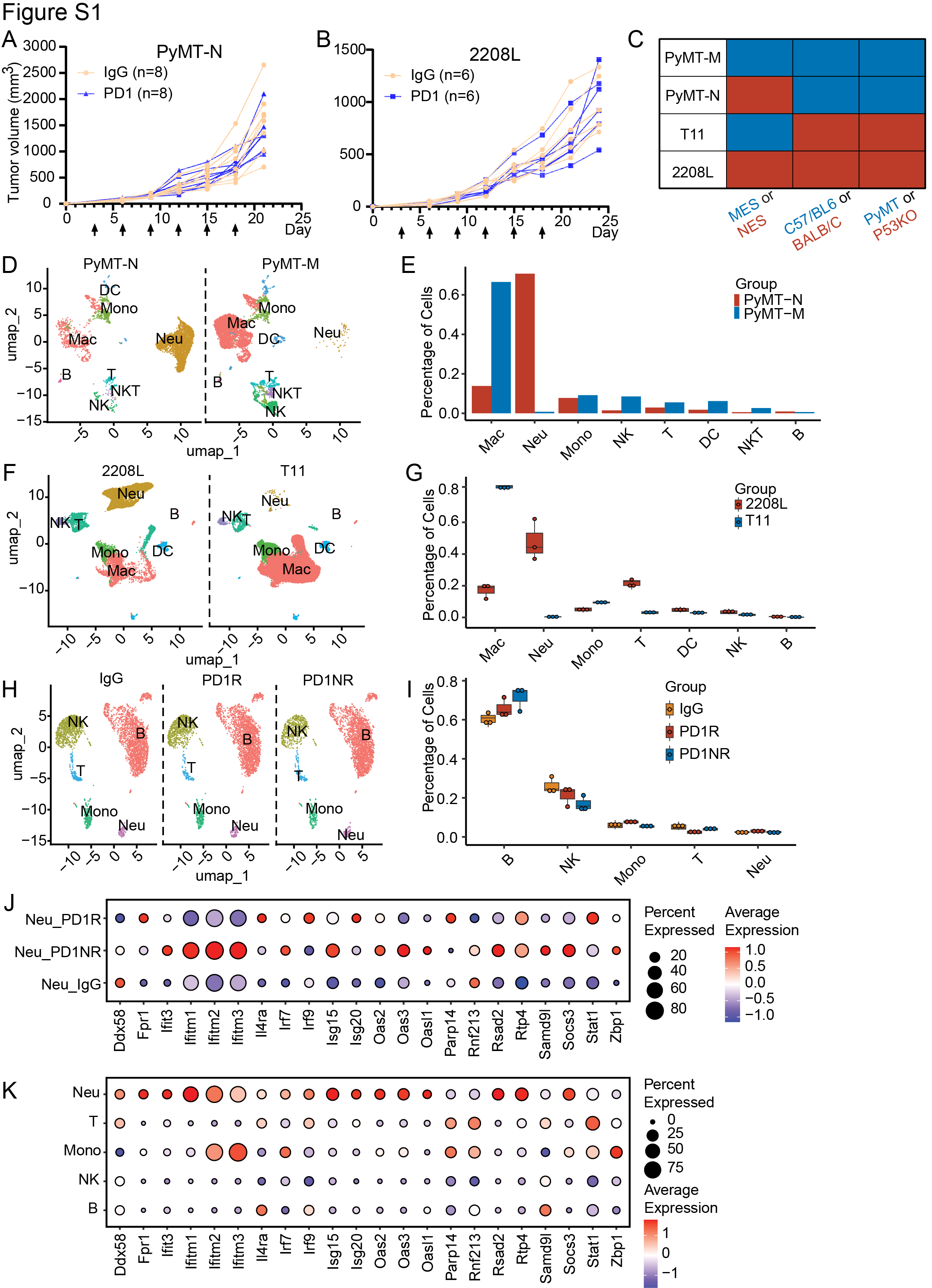

### Supplemental Fig 2

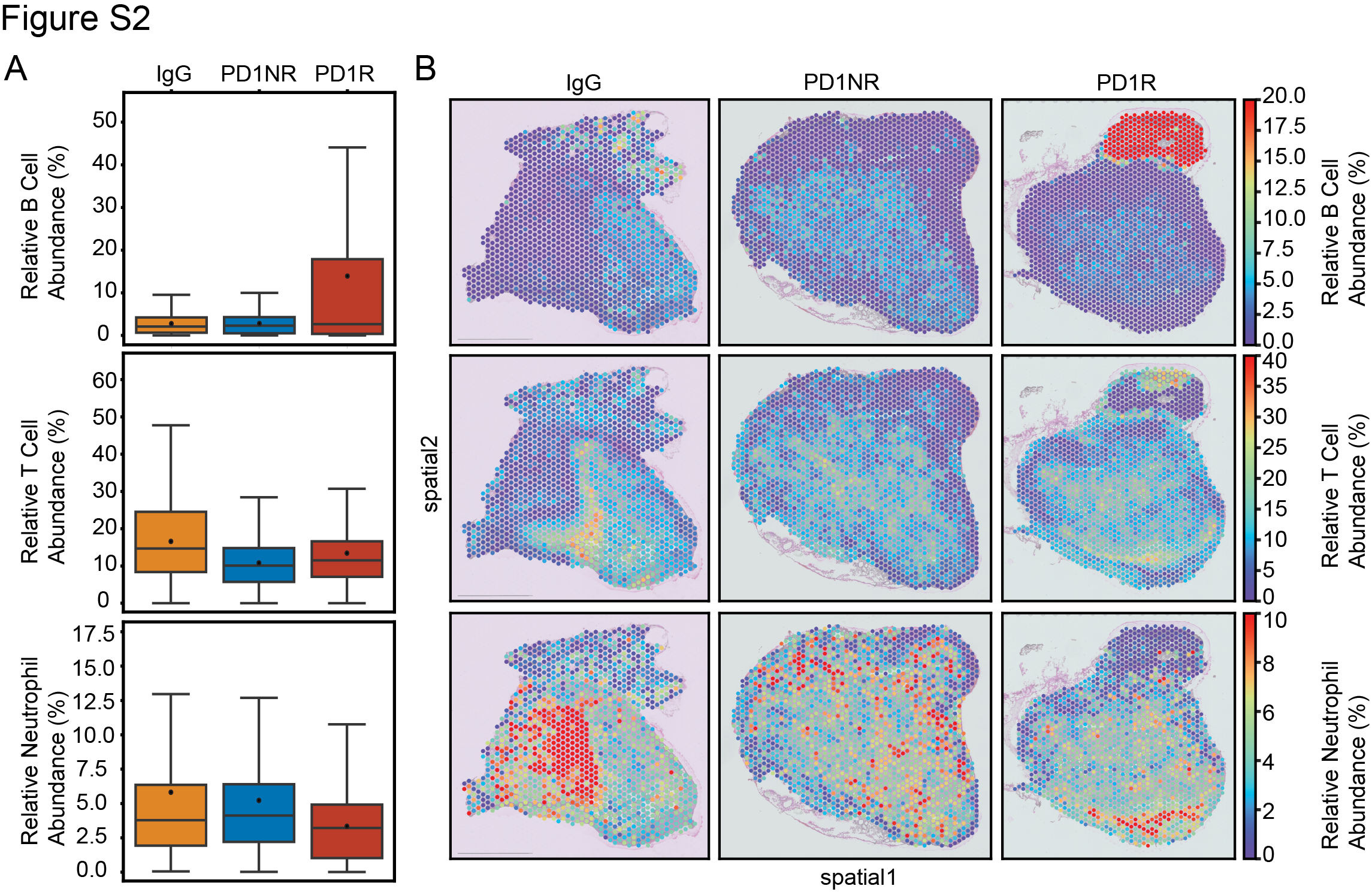

### Supplemental Fig 3

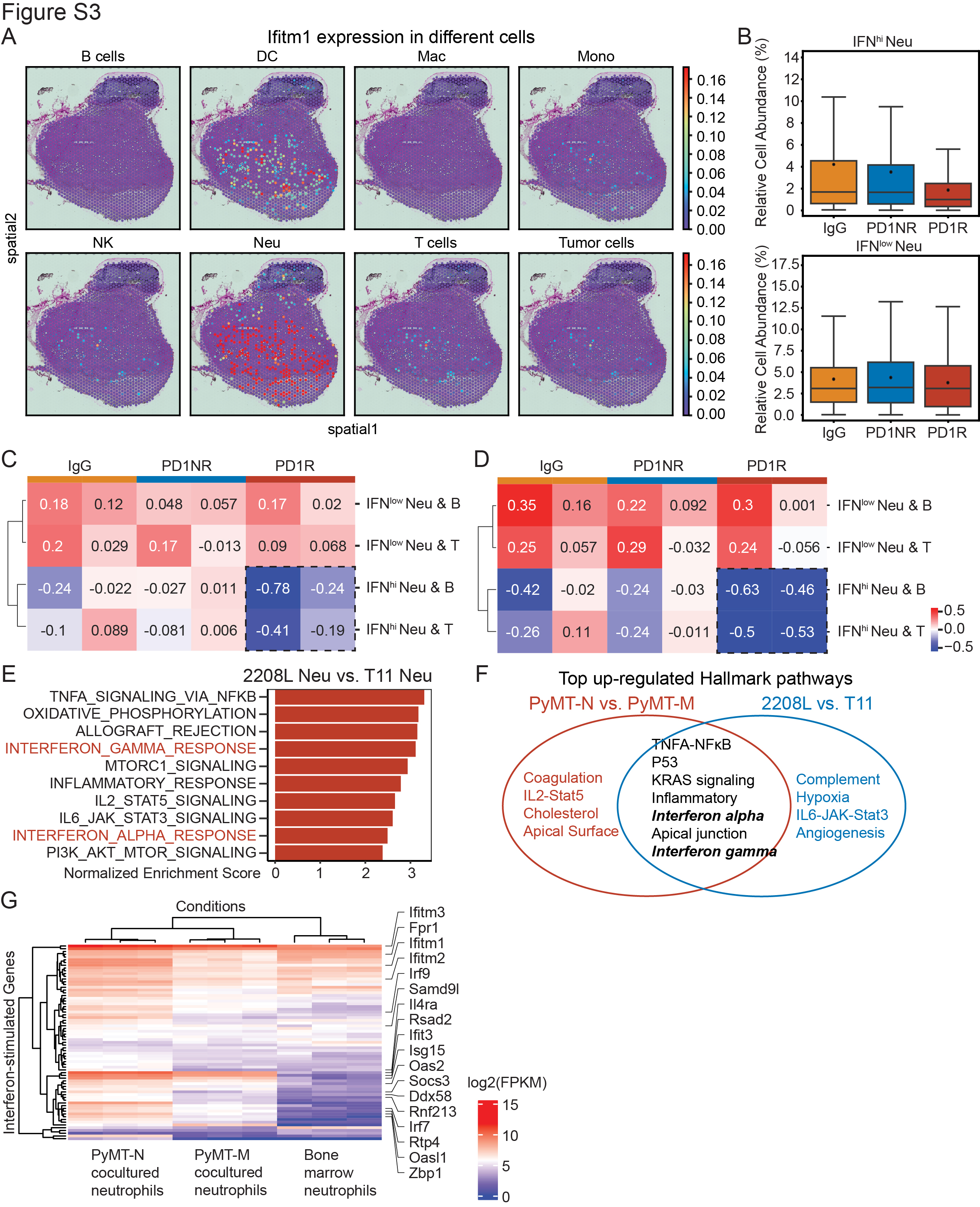

### Supplemental Fig 4

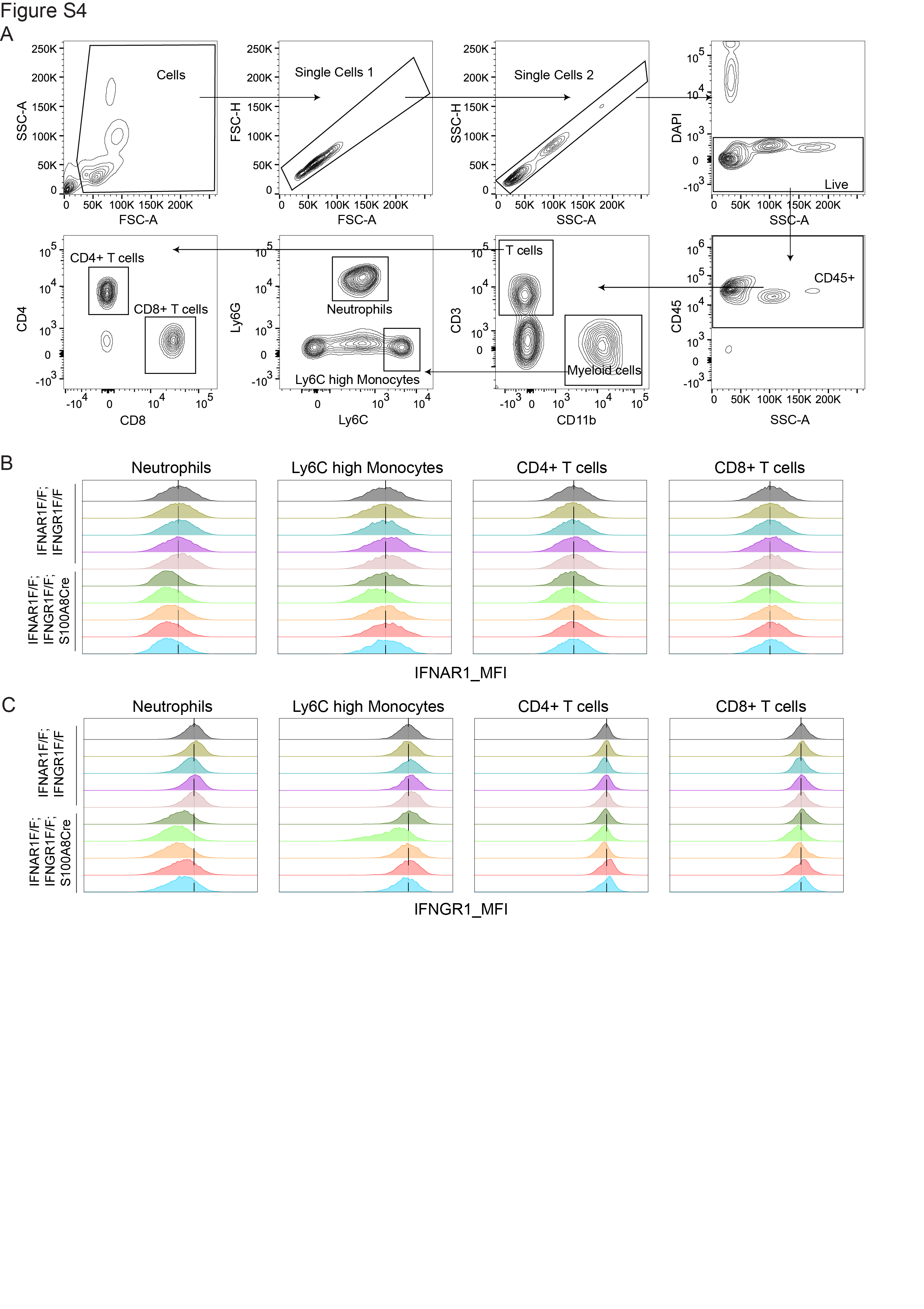

### Supplemental Fig 5

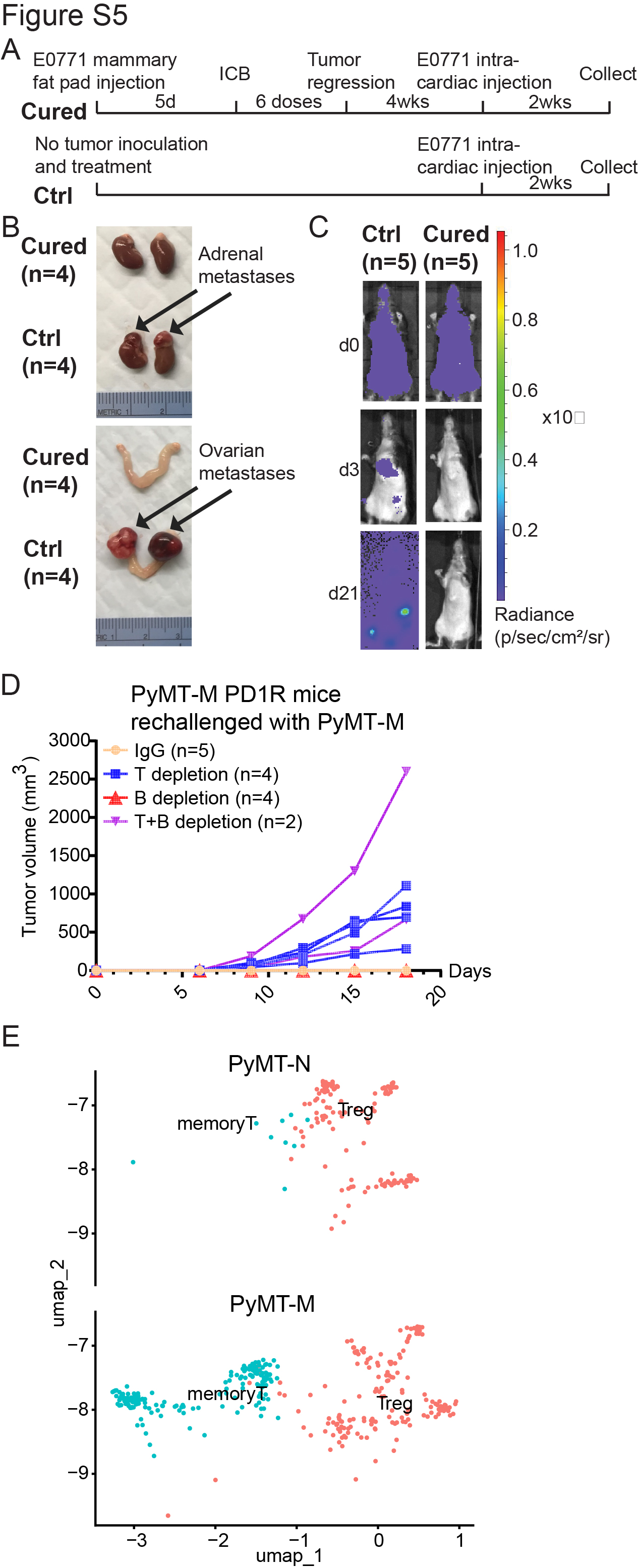
